# Regulation of immune receptor kinases plasma membrane nanoscale landscape by a plant peptide hormone and its receptors

**DOI:** 10.1101/2020.07.20.212233

**Authors:** Julien Gronnier, Christina M. Franck, Martin Stegmann, Thomas A. DeFalco, Alicia Abarca, Michelle Von Arx, Kai Dünser, Wenwei Lin, Zhenbiao Yang, Jürgen Kleine-Vehn, Christoph Ringli, Cyril Zipfel

## Abstract

Spatial partitioning is a propensity of biological systems orchestrating cell activities in space and time. The dynamic regulation of plasma membrane nano-environments has recently emerged as a key fundamental aspect of plant signaling, but the molecular components governing it are still mostly unclear. The receptor kinase FERONIA (FER) controls complex formation of the immune receptor kinase FLAGELLIN SENSING 2 (FLS2) with its co-receptor BRASSINOSTEROID INSENSITIVE 1-ASSOCIATED KINASE 1 (BAK1), and this function is inhibited by the FER ligand RAPID ALKALANIZATION FACTOR 23 (RALF23). Here, we show that FER regulates the plasma membrane nanoscale organization of FLS2 and BAK1. Our study demonstrates that akin to FER, leucine-rich repeat (LRR) extensin (LRXs) proteins contribute to RALF23 responsiveness, regulate BAK1 nanoscale organization and immune signaling. Furthermore, RALF23 perception leads to rapid modulation of FLS2 and BAK1 nanoscale organization and its inhibitory activity on immune signaling relies on FER kinase activity. Our results suggest that perception of RALF peptides by FER and LRXs actively modulates the plasma membrane nanoscale landscape to regulate cell surface signaling by other receptor kinases.

## INTRODUCTION

Multicellular organisms evolved sophisticated surveillance systems to monitor changes in their environment. In plants, receptor kinases (RKs) and receptor proteins (RPs) are the main ligand-binding cell-surface receptors perceiving self, non-self, and modified-self molecules (Hohmann, Lau and Hothorn, 2017). For example, recognition of pathogen-associated molecular patterns (PAMPs) by pattern recognition receptors (PRRs) initiates signaling events leading to PRR-triggered immunity (PTI) (Couto and Zipfel, 2016; Yu *et al*., 2017). The *Arabidopsis thaliana* (hereafter Arabidopsis) leucine-rich repeat receptor kinases (LRR-RKs) FLAGELLIN SENSING 2 (FLS2) and EF-TU RECEPTOR (EFR) recognize the bacterial PAMPs flagellin (or its derived epitope flg22) and elongation factor-Tu (or its derived epitope elf18), respectively (Gómez-Gómez and Boller, 2000; Zipfel *et al*., 2006). Both FLS2 and EFR form ligand-induced complexes with the co-receptor LRR-RK BRASSINOSTEROID INSENSITIVE 1-ASSOCIATED KINASE 1/SOMATIC EMBRYOGENESIS RECEPTOR KINASE 3 (BAK1/SERK3, hereafter BAK1) to initiate immune signaling, such as for instance, the production of apoplastic reactive oxygen species (ROS) (Chinchilla *et al*., 2007; Schulze *et al*., 2010; Schwessinger *et al*., 2011; Sun *et al*., 2013).

We previously showed that the *Catharanthus roseus* RECEPTOR-LIKE PROTEIN KINASE 1-LIKE (CrRLK1L) FERONIA (FER) and the LORELEI-LIKE-GPI ANCHORED PROTEIN 1 (LLG1) are required for flg22-induced FLS2-BAK1 complex formation (Stegmann *et al*., 2017; Xiao *et al*., 2019). Notably, the endogenous RAPID ALKALINIZATION FACTOR 23 (RALF23) is perceived by a LLG1-FER complex, which leads to inhibition of flg22-induced FLS2-BAK1 complex formation (Stegmann *et al*., 2017; Xiao *et al*., 2019). As such, although FER and LLG1 are positive regulator of PTI, RALF23 is a negative regulator. How these components regulate FLS2-BAK1 complex formation remains however unclear.

Members of the CrLKL1s family are involved in RALF perception (Ge *et al*., 2017; Gonneau *et al*., 2018; Liu *et al*., 2021). Among them, FER plays a pivotal role in the perception of several Arabidopsis RALF peptides (Haruta *et al*., 2014; Stegmann *et al*., 2017; Gonneau *et al*., 2018; Zhao *et al*., 2018; Abarca, Franck and Zipfel, 2021; Liu *et al*., 2021). In addition, cell-wall associated LEUCINE RICH REPEAT-EXTENSINs (LRXs) proteins are also involved in CrRLK1L-regulated pathways and were shown to be high-affinity RALF-binding proteins (Mecchia *et al*., 2017; Zhao *et al*., 2018; Dünser *et al*., 2019; Moussu *et al*., 2020). Structural and biochemical analyses have shown that RALF-binding by CrRLK1L/LLGs complexes and LRXs are mutually exclusive and mechanistically distinct from each other (Xiao *et al*., 2019; Moussu *et al*., 2020). While CrRLK1Ls and LRXs have emerged as important RALF-regulated signaling modules, it is still unknown whether LRXs are also involved in RALF23-mediated regulation of immune signaling.

Plasma membrane lipids and proteins dynamically organize into diverse membrane domains giving rise to fluid molecular patchworks (Gronnier *et al*., 2018; Jaillais and Ott, 2020). These domains are proposed to provide dedicated biochemical and biophysical environments to ensure acute, specific and robust signaling events (Gronnier *et al*., 2019; Jacobson, Liu and Lagerholm, 2019). For instance, FLS2 localizes in discrete and static structures proposed to specify immune signaling (Bücherl *et al*., 2017). The cell wall is thought to impose physical constraints on the plasma membrane, limiting the diffusion of its constituents (Feraru *et al*., 2011; Martinière *et al*., 2012). Indeed, alteration of cell wall integrity leads to aberrant protein motions at the plasma membrane (Martinière *et al*., 2012; McKenna *et al*., 2019). Notably, perturbation of the cell wall affects FLS2 nanoscale organization (McKenna *et al*., 2019). Despite its imminent importance, it remains largely unknown how the cell wall and its integrity modulate the organization of the plasma membrane. Interestingly, both CrRLK1Ls and LRXs are proposed cell wall integrity sensors and conserved modules regulating growth, reproduction and immunity (Franck, Westermann and Boisson-Dernier, 2018; Herger *et al*., 2019). However, their mode of action and potential link to RALF perception, are still poorly understood.

Here, we show that FER regulates the plasma membrane nanoscale organization of FLS2 and BAK1. Similarly, we show that LRXs contribute to RALF23 responsiveness, regulate BAK1 nanoscale organization and immune signaling. Importantly, our work reveals an unexpected uncoupling of FER and LRXs modes-of-action in growth and immunity. We demonstrate that RALF23 perception leads to rapid modulation of FLS2 and BAK1 nanoscale organization and that its inhibitory activity on immune signaling requires FER kinase activity. We propose that the regulation of the plasma membrane nanoscale organization by RALF23 receptors underscores their role in the formation of protein complexes and initiation of immune signaling.

## RESULTS AND DISCUSSION

### FER regulates membrane nanoscale organization of FLS2 and BAK1

We combined variable angle total internal reflection fluorescence microscopy (VA-TIRFM) and single-particle tracking to analyze the lateral mobility of FLS2-GFP proteins in transgenic Arabidopsis lines. Two lines expressing FLS2-GFP under the control of its native promoter were crossed with two *FER* knock-out alleles, *fer-2* and *fer-4*. In line with previous reports (Bücherl *et al*., 2017; Tran *et al*., 2020), we observed that FLS2-GFP localizes to static foci in the wild-type (WT) (Movie S1). Consistently, FLS2-GFP single particle trajectories exhibit a confined mobility behavior (Fig. S1, Movie S1). Comparative analysis of the diffusion coefficient (D), which describes the diffusion properties of detected single particles, showed that FLS2-GFP is more mobile in *fer* mutants than in WT (Fig. S1, S2 and Movie S1). To analyze FLS2-GFP organization, we reconstructed images using a temporal averaging of FLS2-GFP fluorescence observed across VA-TIRFM time series. Furthermore, individual image sections were subjected to kymograph analysis. Using this approach, we found that FLS2-GFP fluorescence is maintained into well-defined and static structures in WT, while it appears more disperse and more labile in both *fer* mutants (Fig. 1A-B and S2). To substantiate these observations, we used previously established spatial clustering index (SCI) which describes protein lateral organization (Gronnier *et al*., 2017; Tran *et al*., 2020). As expected, SCI of FLS2-GFP was lower in *fer-4* than in WT (Fig. 1C), indicating disturbance in FLS2-GFP lateral organization.

**Fig. 1.**
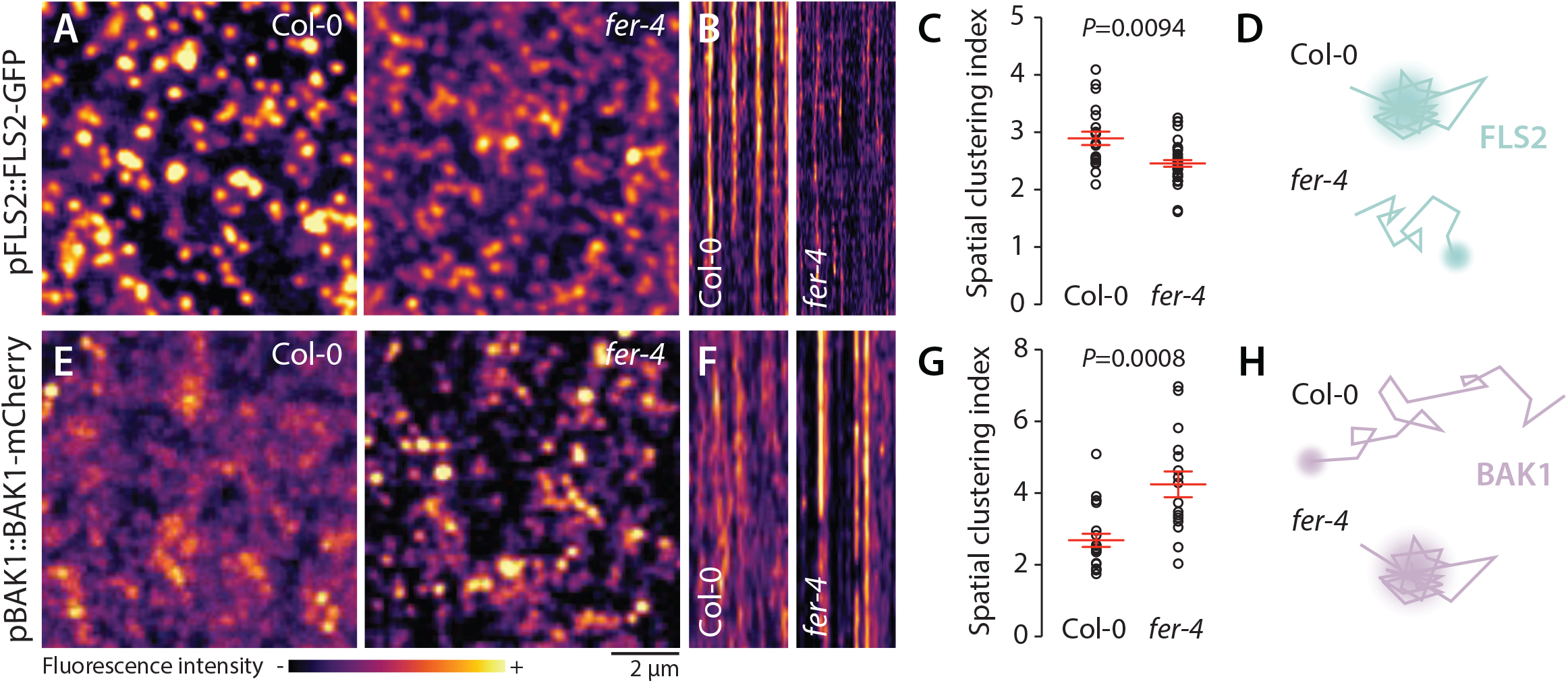
FER regulates the nanoscale organization of FLS2-GFP and BAK1-mCherry. **A and E**. FLS2-GFP and BAK1-mCherry nanodomain organization. Pictures are maximum projection images of FLS2-GFP (**A**) and BAK1-mCherry (**E**) in Col-0 and *fer-4* cotyledon epidermal cells. **B and F**. Representative kymograph showing lateral organization of FLS2-GFP (**B**) and BAK1-mCherry (**F**) overtime in Col-0 and *fer-4*. **C and G**. Quantification of FLS2-GFP (**C**) and BAK1-mCherry (**G**) spatial clustering index. Red crosses and red horizontal lines show mean and SEM, P values reports non-parametric Mann-Whitney test. Similar results were obtained in three independent experiments. **D and H**. Proposed graphical summary of our observations for FLS2-GFP (**D**) and BAK1-mCherry (**H**) nanoscale dynamics.

In *Medicago truncatula* and in yeast, alteration of nanodomain localization has been linked to impaired protein accumulation at the plasma membrane due to increased protein endocytosis (Grossmann *et al*., 2008; Liang *et al*., 2018). To inquire for a potential defect in FLS2 plasma membrane accumulation, we observed sub-cellular localization of FLS2-GFP using confocal microscopy. The analysis revealed a decrease in FLS2-GFP accumulation in *fer* mutants (Fig. S3). Altogether, these results show that *FER* is genetically required to control FLS2-GFP nanoscale organization and accumulation at the plasma membrane.

To further characterize the impact of *FER* loss-of-function in RK organization, we analyzed the behavior of BAK1-mCherry at the plasma membrane. Fluorescence recovery after photobleaching experiments previously suggested that the vast majority of BAK1 molecules are mobile (Hutten *et al*., 2017). Consistent with this result, BAK1-mCherry was much more mobile than FLS2-GFP in the WT background (Sup Movie 2). Given that BAK1 is a common co-receptor for multiple LRR-RK signaling pathways, we hypothesize that BAK1 might dynamically associates with various pre-formed signaling platforms, such as FLS2 nanodomains (Fig. 1, (Bücherl *et al*., 2017)). Under our experimental conditions, we were not able to perform high-quality single particle tracking analysis for BAK1-mCherry (Sup Movie 2, see methods section). However, visual inspection of particles behavior suggests that BAK1-mCherry is less mobile in *fer-4* than in WT (Sup Movie 2). In good agreement, reconstructed VA-TIRFM images and kymographs show that BAK1-mCherry fluorescence is more structured and static in *fer-4* than in WT (Fig. 1F). Furthermore, we observed an increase of BAK1-mCherry SCI in *fer-4* (Fig 1G). Confocal microscopy analysis did not reveal significant differences in BAK1-mCherry plasma membrane accumulation between *fer-4* and WT backgrounds (Fig. S4). Altogether, these data show that loss of *FER* perturbs FLS2 and BAK1 nanoscale organization, albeit in an opposite manner (Fig. 1D and H). Previous reports have similarly shown that altering the composition of the cell wall can lead to opposed effects on the mobility of different proteins. For instance, inhibition of cellulose synthesis increases the mobility of HYPERSENSITIVE INDUCED REACTION 1 (HIR1) (Daněk *et al*., 2020) but limits the mobility of LOW TEMPERATURE INDUCED PROTEIN 6B (Lti6b) (Martinière *et al*., 2012; Daněk *et al*., 2020). Modification of pectin methyl esterification status increases the mobility of FLS2 (McKenna *et al*., 2019) but decreases the mobility of FLOTILIN 2 (FLOT2) (Daněk *et al*., 2020). Collectively, these observations suggest that various membrane environments are differentially regulated by the cell wall and the proposed cell wall integrity sensor FER.

### LRX3, LRX4 and LRX5 regulate BAK1 nanoscale organization and PTI signaling

LRXs are dimeric, cell wall-localized, high-affinity RALF-binding proteins suggested to monitor cell wall integrity in growth and reproduction (Baumberger, Ringli and Keller, 2001; Mecchia *et al*., 2017; Dünser *et al*., 2019; Herger *et al*., 2019, 2020; Moussu *et al*., 2020). Their extensin domain confers cell wall anchoring while their LRR domain mediates RALF binding (Herger *et al*., 2019; Moussu *et al*., 2020). Among the Arabidopsis 11-member *LRX* family, *LRX3, LRX4*, and *LRX5* are the most expressed in vegetative tissues, and the *lrx3 lrx4 lrx5* triple mutant (hereafter *lrx3/4/5*) shows stunted growth and salt hypersensitivity phenotypes reminiscent of the *fer-4* mutant (Zhao *et al*., 2018; Dünser *et al*., 2019). Therefore, we hypothesized that LRXs also regulate immune signaling. Indeed, our co-immunoprecipitation experiments showed that *lrx3/4/5* is defective in flg22-induced FLS2-BAK1 complex formation (Fig. 2A). Consistently, flg22-induced ROS production was reduced in *lrx3/4/5* similar to the levels observed in *fer-4* (Fig. 2B). In addition, we found that *lrx3/4/5* was impaired in elf18-induced ROS production (Fig. 2C), suggesting that, as for FLS2-BAK1 complex formation, *LRX3/4/5* are required for complex formation between EFR and BAK1. Thus, we conclude that LRX3/4/5 are positive regulators of PTI signaling.

**Fig. 2.**
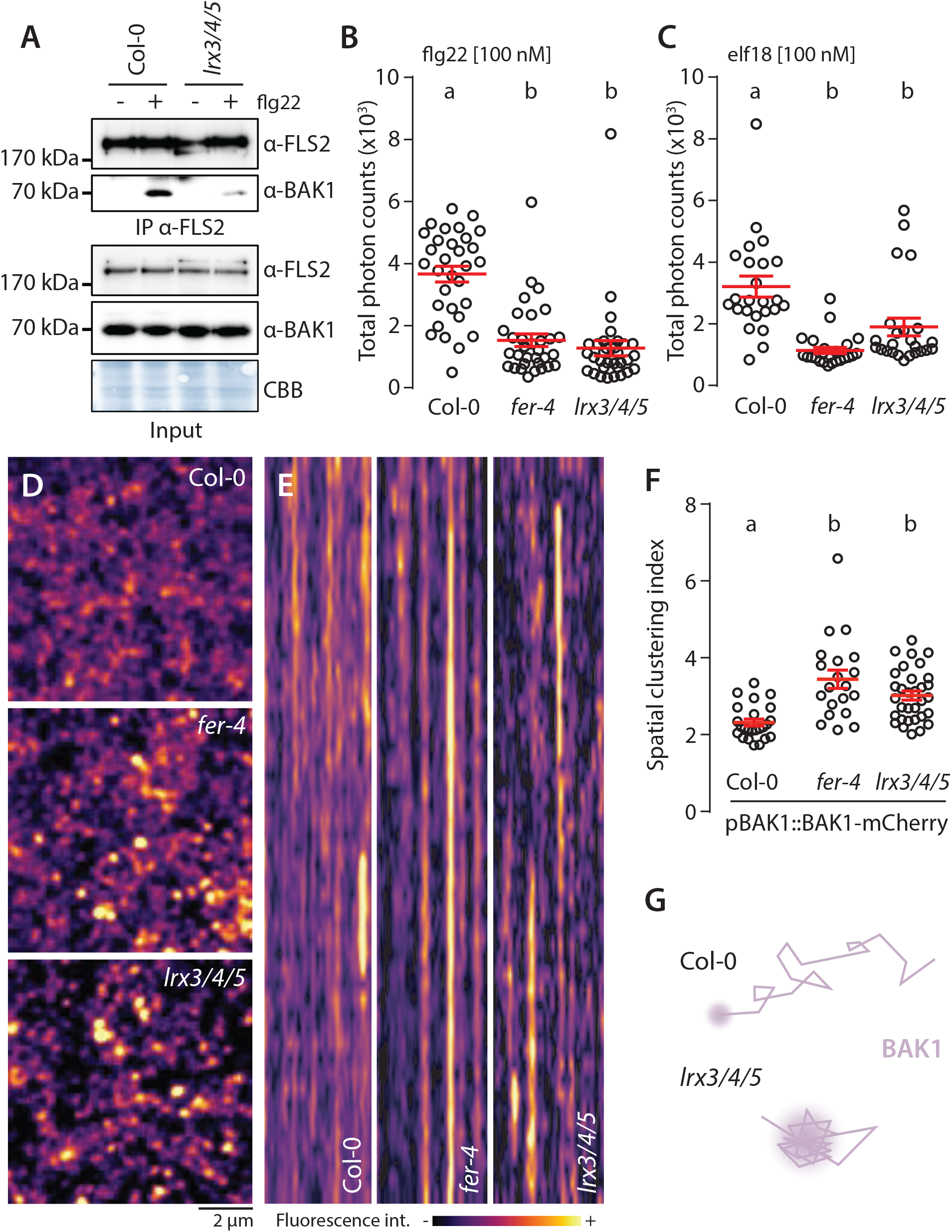
LRX3, LRX4 and LRX5 regulate PTI and BAK1-mCherry organization. **A**. flg22-induced FLS2-BAK1 complex formation. Immunoprecipitation of FLS2 in Arabidopsis Col-0 and *lr3/4/5* seedlings either untreated or treated with 100 nM flg22 for 10 min. Blot stained with CBB is presented to show equal loading. Western blots were probed with α-FLS2, α-BAK1, or α-FER antibodies. Similar results were obtained in at least three independent experiments. **B-C**. ROS production after elicitation with 100 nM elf18 (**B**), or 100 nM flg22 (**C**). Values are means of total photon counts over 40 min. Red crosses and red horizontal lines denote mean and SEM, *n* = 32. Conditions which do not share a letter are significantly different in Dunn’s multiple comparison test (p< 0.0001). **D**. BAK1-mCherry nanodomain organization. Pictures are maximum projection images (20 TIRFM images obtained at 2.5 frames per second) of BAK1-mCherry in Col-0, *fer-4* and *lrx3/4/5* cotyledon epidermal cells. **E**. Representative kymograph showing lateral organization of BAK1-mCherry overtime in Col-0, *fer-4* and *lrx3/4/5*. **F**. Quantification of BAK1-mCherry spatial clustering index, red crosses and red horizontal lines show mean and SEM. Conditions which do not share a letter are significantly different in Dunn’s multiple comparison test (p< 0.001). Similar results were obtained in three independent experiments. **G**. Proposed graphical summary of our observations for BAK1-mCherry nanoscale dynamics in *lrx3/4/5*.

We then asked whether, similar to FER, LRX3/4/5 regulate plasma membrane nanoscale organization. We crossed lines expressing FLS2-GFP and BAK1-mCherry under the control of their respective native promoter with *lrx3/4/5* background. However, despite several attempts, we could not retrieve homozygous *lrx3/4/5* lines expressing FLS2-GFP. Nonetheless, VA-TIRFM and confocal imaging showed that, like in *fer-4*, BAK1-mCherry is more organized and more static in *lrx3/4/5* (Fig. 2D-E, Sup Movie 3), and that BAK1-mCherry plasma membrane localization is not affected by the loss of *LRX3/4/5* (Fig. S4). Thus, like in *fer* mutants, perturbation in PTI signaling observed in *lrx3/4/5* correlates with alterations of plasma membrane RK organization.

Formerly, LRX3, LRX4 and LRX5 have been proposed to sequester RALF peptides to prevent internalization of FER and inhibition of its function (Zhao *et al*., 2018). Following this logic, defects in PTI observed in *lrx3/4/5* could be explained by a depletion of FER at the plasma membrane. However, our confocal microscopy analysis and western blotting with anti-FER antibodies indicated that FER accumulation and plasma membrane localization is not affected in *lrx3/4/5* (Fig. S5). Furthermore, VA-TIRFM revealed that FER-GFP transiently accumulates in dynamic foci, independently of *LRX3/4/5* (Fig. S6, Sup movie 4). Together, these results suggest that LRX3/4/5 do not prevent RALF association with FER to modulate PTI. Moreover, our results suggest that active monitoring by the proposed cell wall integrity sensors FER and LRXs regulates plasma membrane nanoscale dynamics of RKs.

The ability of LRX3/4/5 to associate with RALF23 *in planta* (Zhao *et al*., 2018) prompted us to test if LRX3/4/5 are required for RALF23 responsiveness. Indeed, *LRX3, LRX4* and *LRX5* were required for RALF23-induced inhibition of ROS production upon elf18 treatment (Fig. S7A). Similarly, we observed a decrease in the inhibition of seedlings growth triggered by RALF23 in *lrx3/4/5* compared to WT (Fig. S7B). Altogether, these data show that LRX3/4/5 contribute to RALF23 responsiveness (Fig. S7C), and that LRXs and FER have analogous functions in regulating PTI.

We next asked if FER and LRX3/4/5 form a complex. For this, we made use of a deleted version of LRX4 lacking its extensin domain (LRX4^ΔE^), previously used to assess protein complex formation (Dünser *et al*., 2019; Herger *et al*., 2020). Consistent with previous reports on transient expression assays, (Dünser *et al*., 2019; Herger *et al*., 2020), co-immunoprecipitation experiments with stably transgenic Arabidopsis showed that FER associates constitutively with LRX4^ΔE^-FLAG, and that RALF23 treatment does not modulate this association (Fig. S8). This suggests that FER-LRX complex-mediated direct monitoring of the cell wall (Dünser *et al*., 2019; Herger *et al*., 2019) is not regulated by RALF23. In agreement with structural and biochemical analyses of RALF-binding by CrRLK1L/LLGs and LRXs (Moussu *et al*., 2020), FER-LLG1 and LRX3/4/5 may form distinct RALF23 receptor complexes. As in the cases of pollen tube and root hair growth and integrity (Ge *et al*., 2017; Mecchia *et al*., 2017; Moussu *et al*., 2020; Dünser *et al*., 2019; Herger *et al*., 2020), future investigations are thus needed to understand the exact molecular link between RALF-binding LRXs and CrRLK1s.

### Functional dichotomy of FER and LRXs in regulating growth and immunity

In line with previous reports our data shows that FER and LRXs can form a complex ((Dünser *et al*., 2019; Herger *et al*., 2019), Fig. S8). Moreover, they are known to associate with the cell wall (Baumberger, Ringli and Keller, 2001; Feng *et al*., 2018), and are proposed to relay co-jointly its properties (Dünser *et al*., 2019; Herger *et al*., 2019). We thus asked if direct cell wall sensing underlies FER and LRXs function in PTI. In the context of growth and cell expansion, plants overexpressing LRX4^ΔE^, a truncated version of LRX4, are phenotypically reminiscent of *lrx3/4/5* triple mutants and the *fer-4* single mutant (Dünser *et al*., 2019). This dominant negative effect is proposed to be caused by competition of the overexpressed truncated LRX4^ΔE^ with endogenous LRXs and consequent loss of cell wall anchoring (Dünser *et al*., 2019). Similarly, overexpression of LRX1^ΔE^ inhibits root hair elongation, phenocopying loss of function of *LRX1* and *LRX2* (Herger *et al*., 2020). By contrast, we observed that LRX4^ΔE^ does not affect flg22-induced interaction between FLS2 and BAK1 (Fig. S9A). In good agreement with this notion, overexpression of LRX4^ΔE^ did not affect flg22 nor elf18-induced ROS production (Fig. S9B-C). To corroborate these results, we tested inhibition of root growth triggered by flg22 treatment. Consistent with the positive role of FER and LRX3/4/5 in PTI, we observed that *fer-4* and *lrx3/4/5* are hyposensitive to flg22 treatment (Fig S9D). By contrast, overexpression of LRX4^ΔE^ did not affect inhibition of root growth by flg22 (Fig <u>S9D</u>). In addition, we observed that LRX4^ΔE^ does not impact RALF23 responsiveness (Fig S9E). Altogether, these data suggest that the function of LRX3/4/5 in PTI is distinct from their role during growth.

The ectodomain of FER contains two malectin-like domains, malA and malB (Fig 3A), which share homology with malectin, a carbohydrate-binding protein from *Xenopus laevis* (Boisson-Dernier, Kessler and Grossniklaus, 2011). Despite lacking the canonical carbohydrate-binding site of malectin (Moussu *et al*., 2018; Xiao *et al*., 2019), malA and malB have been proposed to bind pectin *in vitro* (Feng *et al*., 2018) and FER-mediated cell wall-sensing regulates morphogenesis (Duan *et al*., 2010; Lin *et al*., 2018). To investigate if direct cell wall-sensing underlies FER‘s function in regulating PTI, we used transgenic lines expressing a FER truncated mutant lacking the malA-domain C-terminally fused to YFP (FER^ΔmalA^-YFP) in the *fer-4* mutant background (Fig 3B). We observed that FER^ΔmalA^-YFP did not complement cell shape or root hair elongation defect of *fer-4* (Fig. 3C-D), emphasizing the importance of malA in FER-regulated cell morphogenesis. In contrast, immuno-precipitation assays showed that FER^ΔmalA^-YFP fully complements flg22-induced complex formation between endogenous FLS2 and BAK1 (Fig. 3E) as well as ROS production in response to flg22 and elf18 (Fig. 3F-G). Altogether, these data suggest that malA-mediated cell wall sensing underlies specific function(s) of FER in regulating growth and cell morphology, but is dispensable for FER’s role in PTI. Interestingly, we observed that expression of FER^ΔmalA^-YFP restores inhibition of growth triggered by RALF23, suggesting that malB is sufficient for RALF responsiveness (Fig. 3H), as suggested by its physical interaction with RALF23 (Xiao *et al*., 2019). While we cannot formally exclude the implication of pectin-binding by malB in regulating immunity, the contrasted context-dependent functionality of FER^ΔmalA^-YFP suggests that FER’s function in PTI is primarily mediated by RALF perception. Altogether, our data indicate molecular and functional dichotomy of FER and LRXs in regulating growth and immunity.

**Fig. 3.**
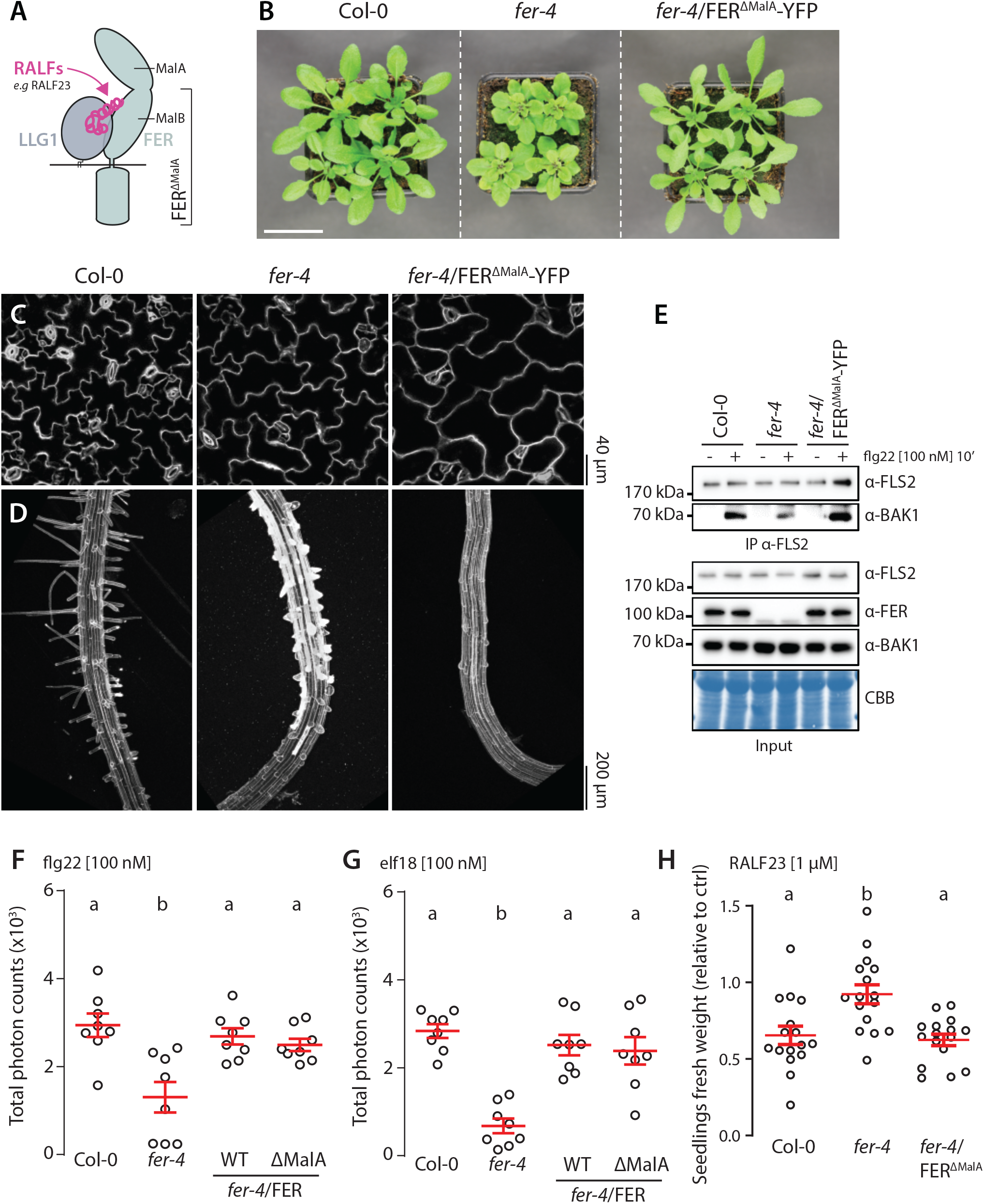
FER malectin A domain regulates cell morphogenesis not PTI. **A**. Proposed graphical representation of RALF23 perception by FER-LLG1 complex. **B**. Morphology of 4-week-old Arabidopsis plants, scale bar indicates 5 cm. **C-D**. Confocal microscopy pictures of 5-day-old seedlings cotyledon (**C**) and root (**D**) stained with propidium iodide. Similar results were obtained in at least three independent experiments. **E**. flg22-induced FLS2-BAK1 complex formation. Immunoprecipitation of FLS2 in Arabidopsis Col-0, *fer-4*, and *fer-4*/p35S::FER^ΔMalA^-YFP seedlings that were either untreated or treated with 100 nM flg22 for 10 min. Blot stained with Coomassie brilliant blue (CBB) is presented to show equal loading. Western blots were probed with α-FLS2, α-BAK1, or α-FER antibodies. Similar results were obtained in at least three independent experiments. **F-G**. ROS production after elicitation with 100 nM flg22 (**F**), or 100 nM elf18 (**G**). Values are means of total photon counts over 40 min, *n* = 8. Red crosses and red horizontal lines denote mean and SEM, respectively. Conditions which do not share a letter are significantly different in Dunn’s multiple comparison test (p< 0.0001). **H**. Fresh weight of 12-day-old seedlings grown in the absence (mock) or presence of 1 µM of RALF23 peptide. Fresh weight is expressed as relative to the control mock condition. Similar results were obtained in at least three independent experiments. Conditions which do not share a letter are significantly different in Dunn’s multiple comparison test (p< 0.001).

**Fig. 4.**
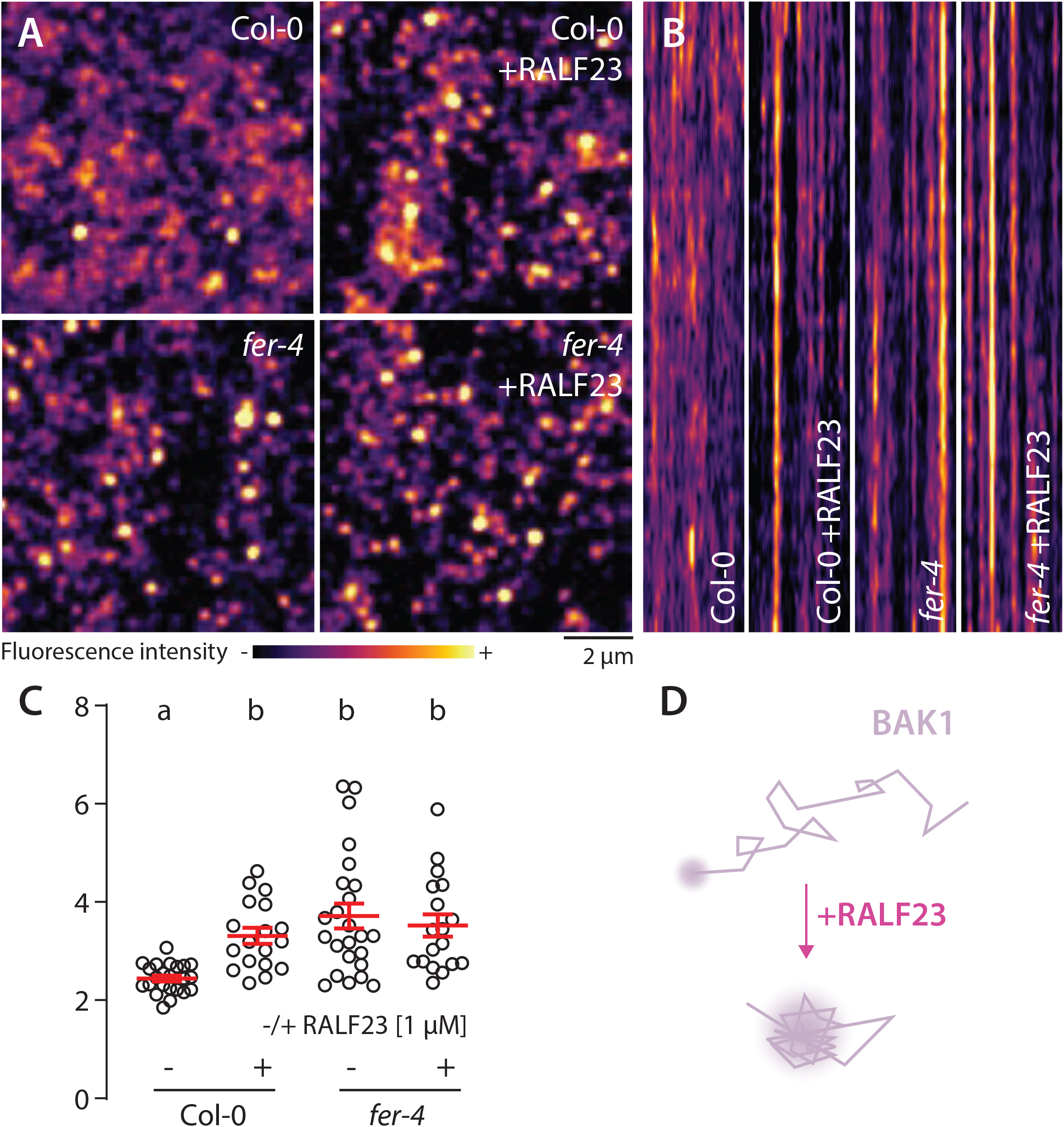
RALF23 perception regulates BAK1-mCherry organization. **A**. BAK1-mCherry nanodomain organization (pBAK1::BAK1-mCherry). Pictures are maximum projection images of BAK1-mCherry in Col-0 and *fer-4* cotyledon epidermal cells with or without 1 µM RALF23 treatment (2 to 30 min). **B**. Representative kymograph showing lateral organization of BAK1-mCherry overtime in Col-0 and *fer-4* with or without 1 µM RALF23 treatment. **C**. Quantification of BAK1-mCherry spatial clustering index, red crosses and red horizontal lines show mean and SEM. Conditions which do not share a letter are significantly different in Dunn’s multiple comparison test (p< 0.001). Similar results were obtained in three independent experiments. **D**. Proposed graphical summary of our observations for BAK1-mCherry nanoscale dynamics upon RALF23 treatment.

### RALF23 alters FLS2 and BAK1 organization and function through active FER signaling

We next asked if RALF23 activity is mediated by active FER signaling. We used a kinase-dead mutant, FER^K565R^ C-terminally fused to GFP, expressed in *fer* knock-out backgrounds and selected lines accumulating it to level similar to endogenous FER in WT background (Fig. S10; Fig S11 (Chakravorty, Yu and Assmann, 2018)). Interestingly, we observed that FER^K565R^-GFP complements *fer*’s defect in FLS2-BAK1 complex formation (Fig. S10A) and PAMP-induced ROS production (Fig. S10B-C). In contrast, we observed that inhibition of FLS2-BAK1 complex formation by RALF23 depends on FER kinase activity (Fig. S11B. Similarly, inhibition of elf18-induced ROS production and seedlings growth inhibition by RALF23 depended on FER kinase activity (Fig. S11C). Overall, these data show that inhibition by RALF23 is mediated by active FER signaling while FER’s positive role in immune signaling is kinase activity-independent. We next asked if inhibition of FLS2-BAK1 complex formation by RALF23 correlates with a modulation of FLS2 or BAK1 nanoscale organization. VA-TIRFM imaging showed an increase of FLS2-GFP mobility and an alteration of FLS2-GFP nanodomain organization within minutes of RALF23 treatment (Fig. S12 and S13, Movie S7, imaging performed 2 to 30 min post-treatment Fig. S14). Similarly, we observed that RALF23 treatment affects BAK1-mCherry nanoscale organization (Fig. 5, Fig. S14, Movie S2). These data suggest that RALF23 perception leads to rapid modification of FLS2 and BAK1 membrane organization and thereby potentially inhibits their association. It will be important in the future to identify the components mediating RALF23 signaling and modification of FLS2 and BAK1 nanoscale dynamics. In sum, our study unravels the regulation of FLS2 and BAK1 nanoscale organization by the RALF receptors FER and LRX3/4/5. The function of RALF receptors in other processes might similarly rely on the regulation of RK nanoscale dynamics, and the identification of the corresponding regulated RKs is an exciting prospect for future investigation.

## ACKNOWLEDGMENTS

We thank all present and past members of the Zipfel laboratory for fruitful discussions and comments on the manuscript. We thank the members of the Grossniklaus, Ringli, Sanchez-Rodriguez and Keller laboratories for sharing results and comments during our stimulating CCWI meetings. We thank Vera Gorelova, Yvon Jaillais and Birgirt Kemmerling for comments on the manuscript. This research was funded by the Gatsby Charitable Foundation (C.Z.), the University of Zürich (C.Z.), the European Research Council under the Grant Agreements 309858 and 773153 (grants PHOSPHinnATE and IMMUNO-PEPTALK to C.Z.) and 639678 (grant AuxinER to J.K-V), the Swiss National Science Foundation (grant no. 31003A_182625 to C.Z. and 31003A_166577/1 to C.R.), and the Austrian science fund (FWF; P 33044 to J.K-V). J.G., C.M.F. and T.A.D. were supported by Long-Term Fellowships from the European Molecular Biology Organization (EMBO) (numbers 438-2018, 512-2019 and 100-2017, respectively), while M.S. was supported by a post-doctoral fellowship (STE 2448/1) from the Deutsche Forschungsgemeinschaft (DFG) and K.D. by a doctoral fellowship from the Austrian Academy of Sciences (ÖAW). We thank Dr. Sarah Assman for providing segregating lines of *fer-4*/pFER::FER^K565R^-GFP, Prof. Sacco de Vries for kindly providing Col-0/pBAK1::BAK1-mCherry line and Dr. Silke Robatzek for Col-0/pFLS2::FLS2-GFP lines, and Dr. Nana Keinath for kindly offering *fer-2*/pFER::FER^K565R^-GFP lines.

## FIGURE LEGENDS

**Fig. S1.**
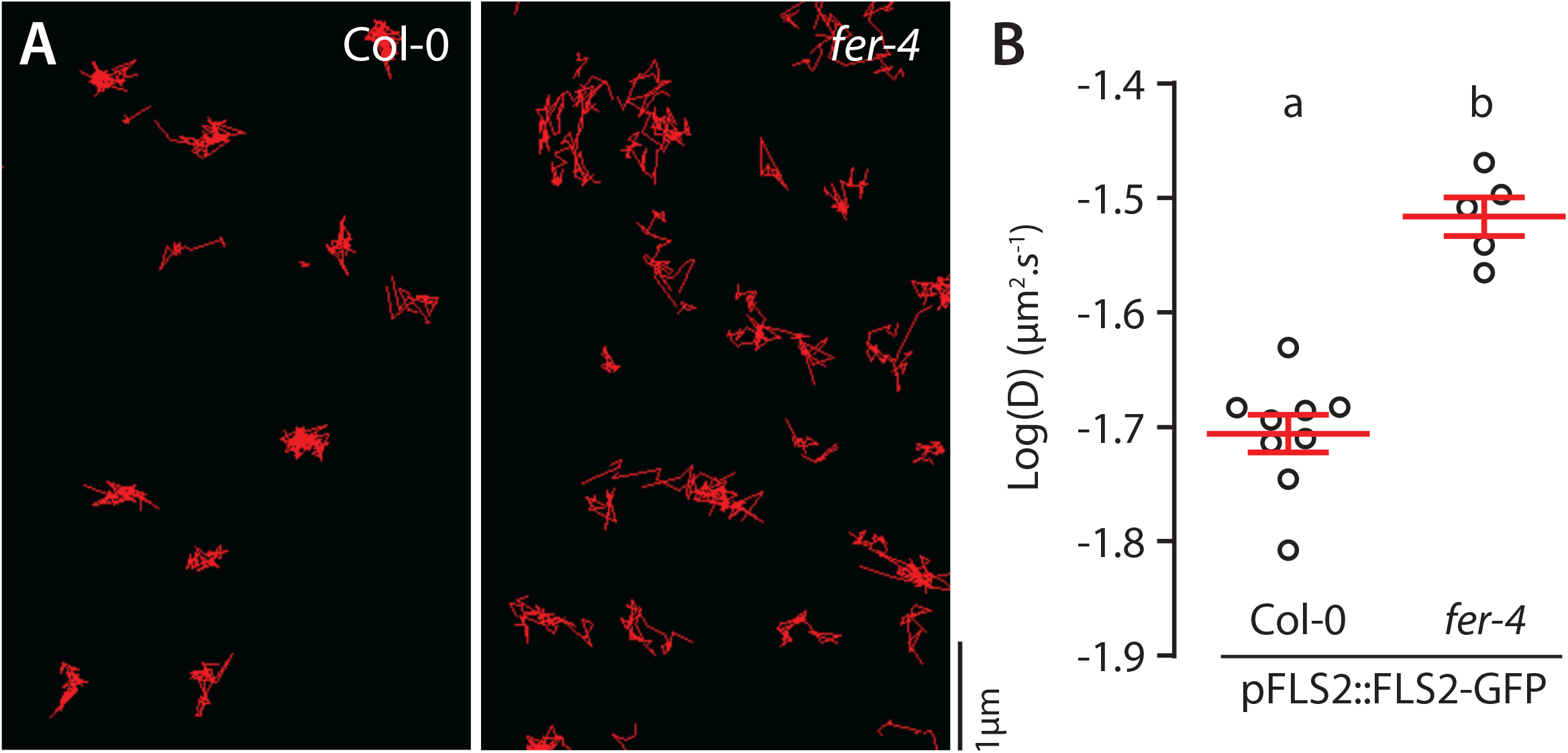
Analysis of FLS2-GFP single particle dynamics in *fer-4*. **A**. Representative images of FLS2-GFP single-particles tracked in Col-0 and *fer-4* cotyledon epidermal cells of 5-day-old seedlings. **B**. Quantification of FLS2-GFP diffusion coefficient (D). Similar results were obtained in three independent experiments.

**Fig. S2.**
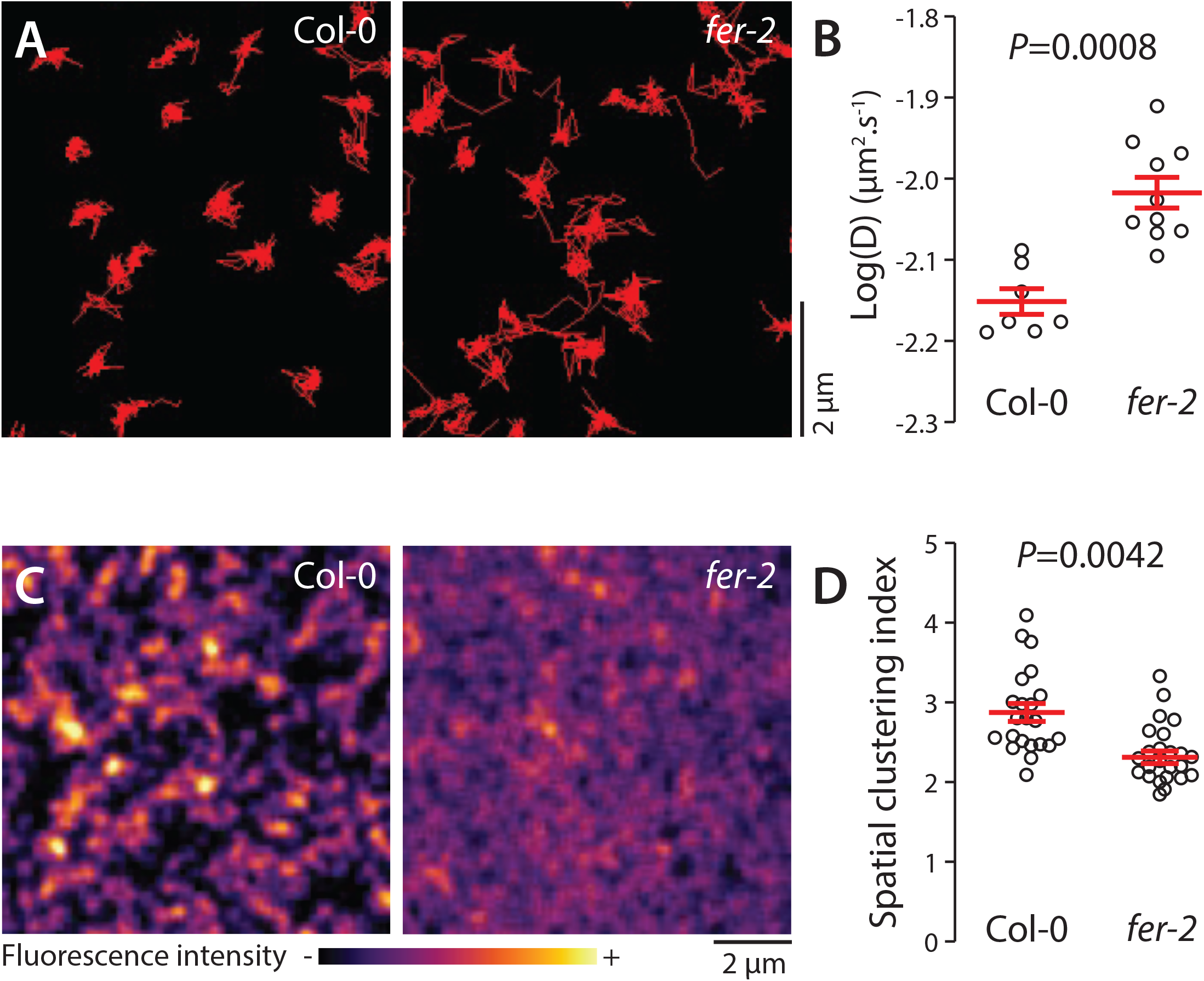
Analysis of FLS2-GFP organization and dynamics in *fer-2*. **A**. Representative images of FLS2-GFP single-particles tracked in Col-0 and *fer-2* cotyledon epidermal cells of 5-day-old seedlings. **B**. Quantification of FLS2-GFP diffusion coefficient (D). Similar results were obtained in three independent experiments. **C**. Pictures are maximum projection images of FLS2-GFP in Col-0 and *fer-2* cotyledon epidermal cells. **D**. Quantification of FLS2-GFP spatial clustering index in Col-0 and *fer-2*. Red crosses and red horizontal lines show mean and SEM, P values reports non-parametric Mann-Whitney test. Similar results were obtained in three independent experiments.

**Fig. S3.**
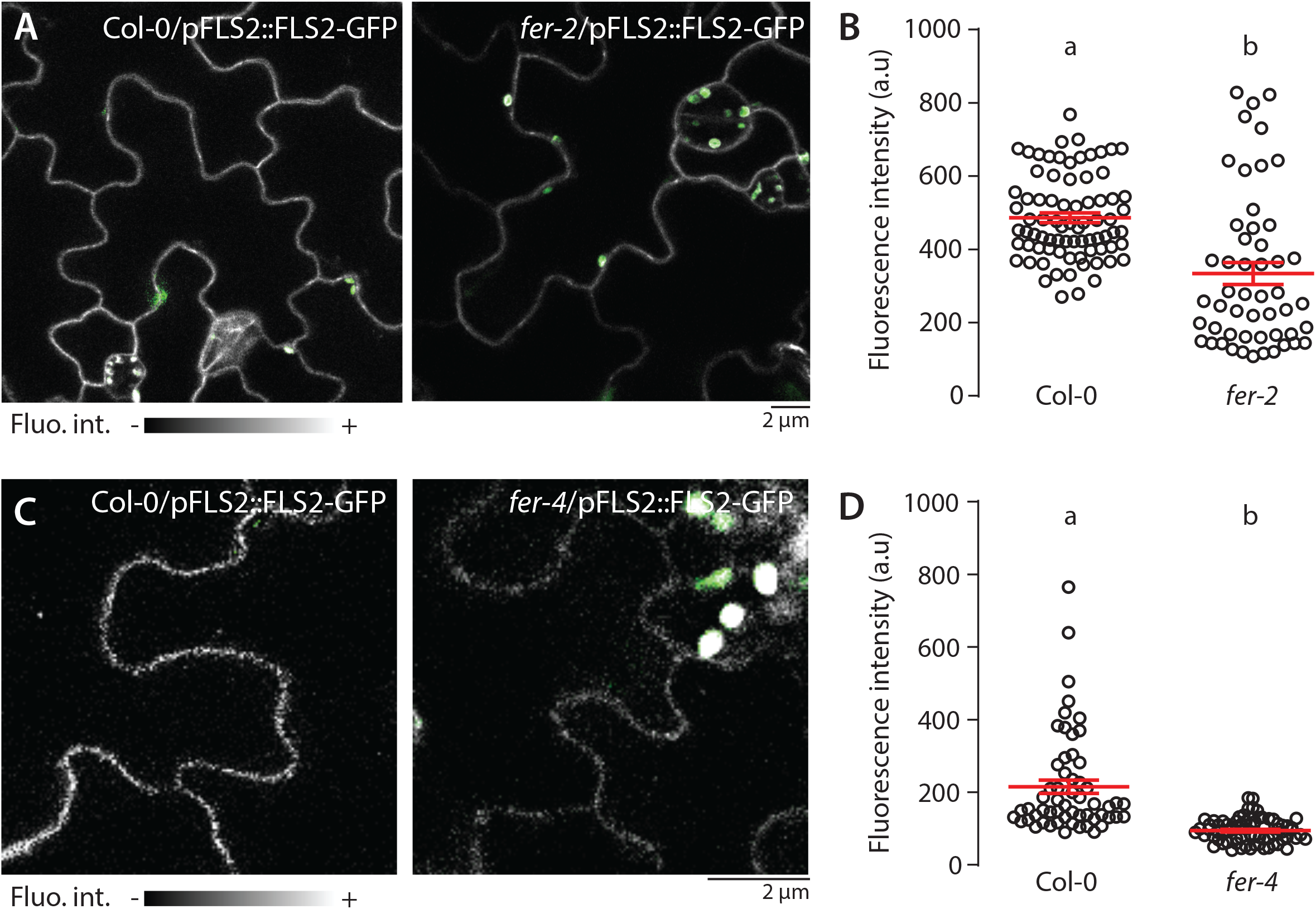
FLS2-GFP accumulation at the PM is altered in *fer* mutants. **A-D**. Representative confocal microscopy pictures of FLS2-GFP driven by its native promoter in Col-0 and *fer-2* (**A**) and *fer-4* (**C**) cotyledon epidermal cells of 5-day-old seedlings. Scale bar indicate 2 µm. Quantification of FLS2-GFP fluorescence intensity at the plasma membrane in Col-0 and *fer-2* (**B**) and *fer-4* (**D**) cotyledon epidermal cells of 5-day-old seedlings. Two-way student’s t test. Similar results were obtained in at least three independent experiments.

**Fig. S4.**
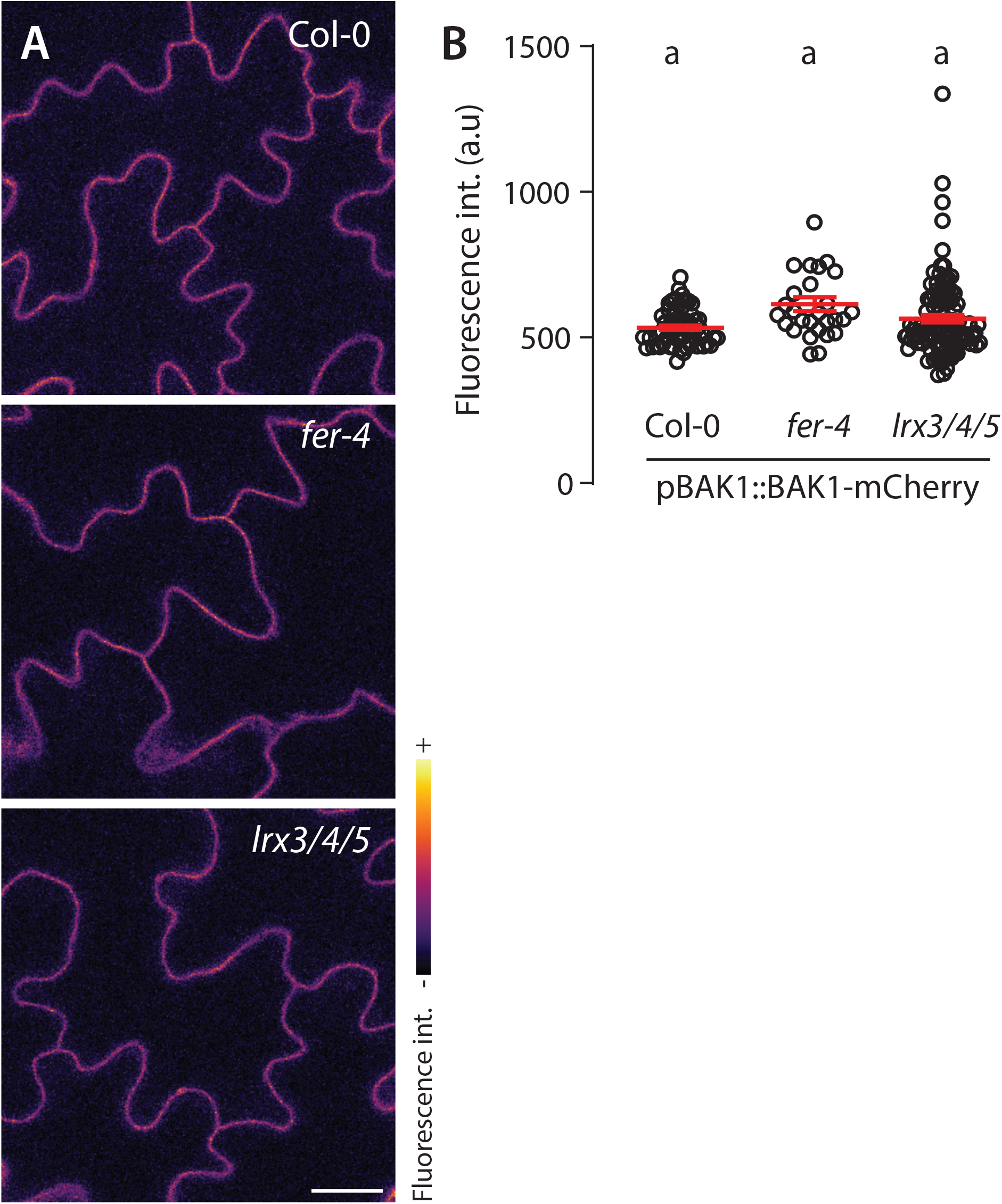
Subcellular localization of BAK1-mCherry in *fer-4* and *lrx3/4/5*. **A**. Representative confocal microscopy pictures of BAK1-mCherry driven by its native promoter in Col-0, *fer-4* and *lrx3/4/5* cotyledon epidermal cells of 5-day-old seedlings. Scale bar indicates 20 µm. **B**. Quantification of BAK1-mCherry fluorescence intensity at the plasma membrane. Letters indicate a Dunn’s multiple comparison statistical test.

**Fig. S5.**
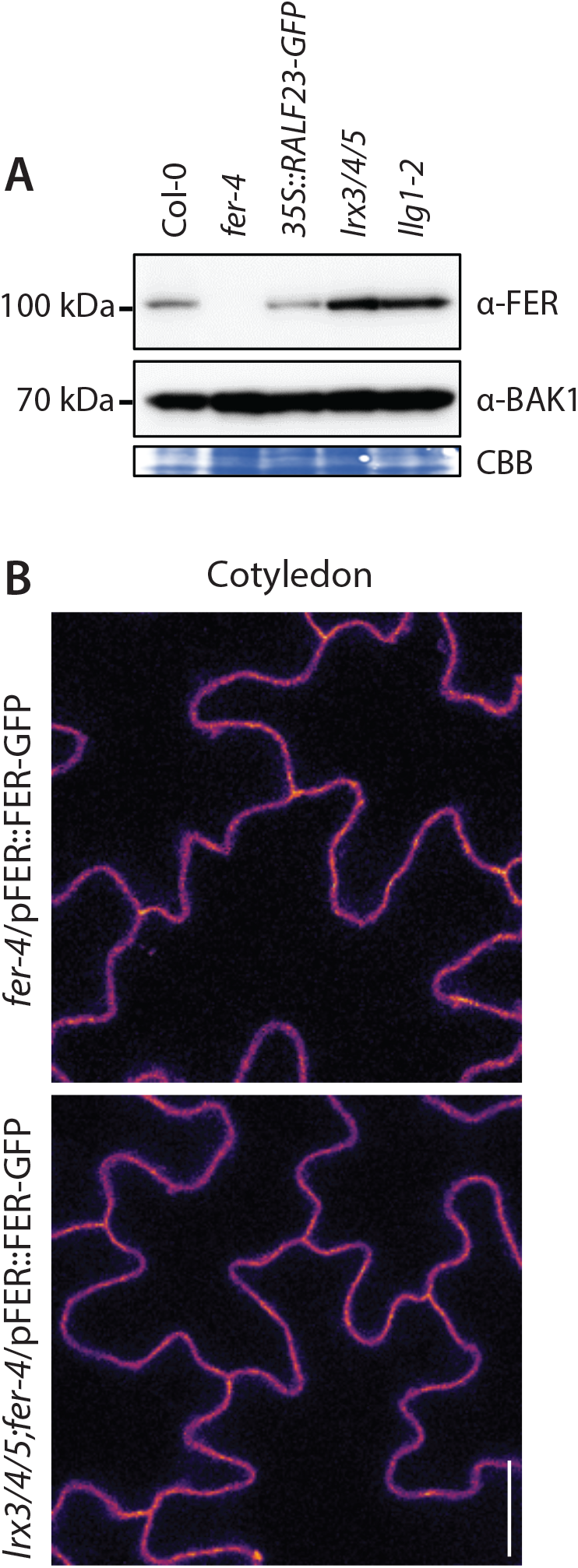
*LRX3, LRX4* and *LRX5* are dispensable for FER plasma membrane localization and accumulation. **A**. Accumulation of endogenous FER detected by western blot. Total proteins were extracted from 2-week-old seedlings. Blot stained with CBB is presented to show equal loading. **B-C**. Representative confocal microscopy pictures of FER-GFP driven by its native promoter in *fer-4* and *lrx3/4/5* cotyledon epidermal cells (**B**) and root hair (**C**) of 5-day-old seedlings. Scale bar indicates 20 µm.

**Fig. S6.**
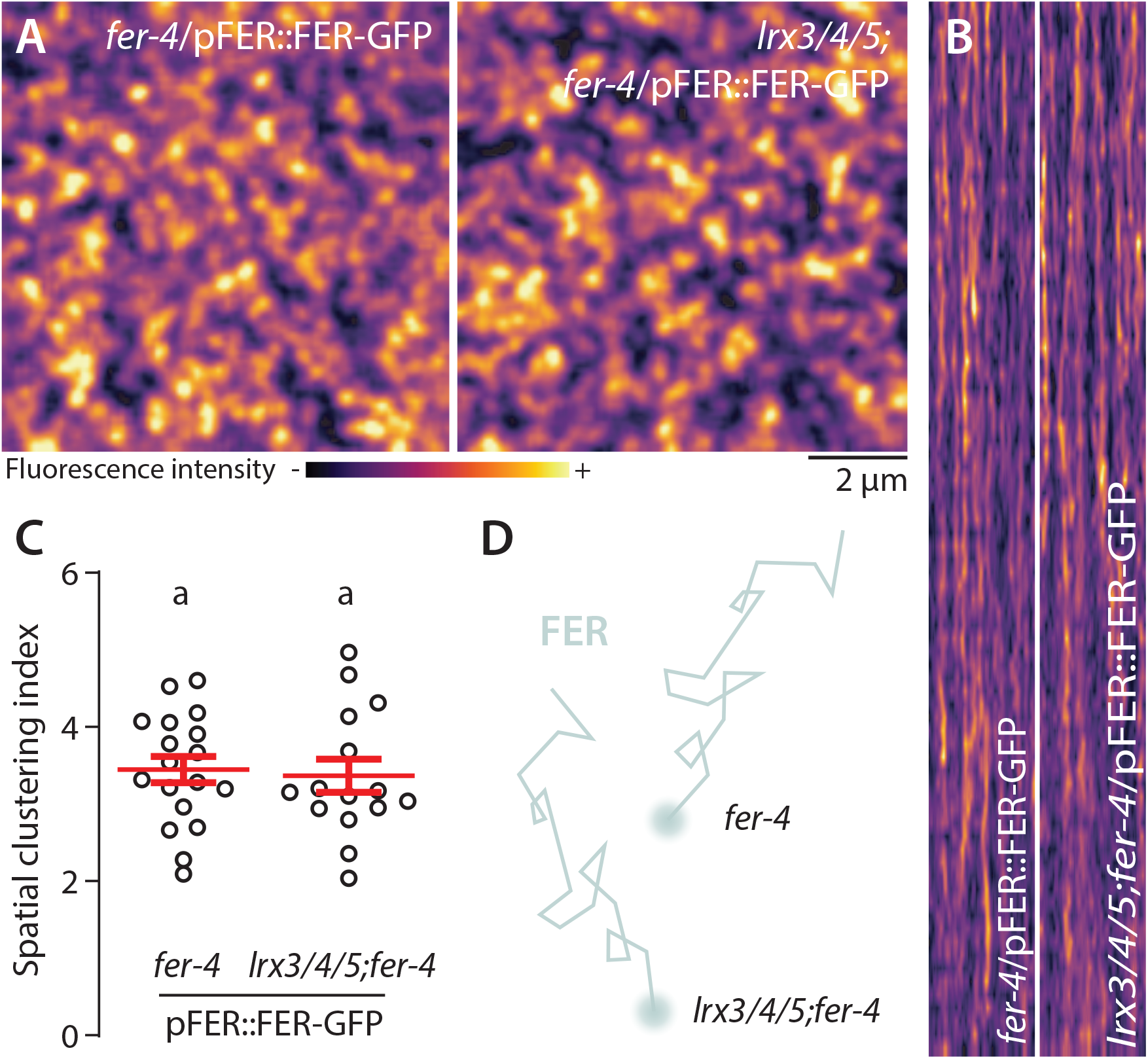
*LRX3, LRX4* and *LRX5* are dispensable for FER-GFP nanoscale organization. **A**. FER-GFP nanodomain organization. Pictures are maximum projection images (20 TIRFM images obtained at 5 frames per second) of FER-GFP in *fer-4* and *lrx3/4/5* cotyledon epidermal cells. **B**. Representative kymograph showing lateral organization of FER-GFP overtime in *fer-4* and *lrx3/4/5*. **C**. Quantification of FER-GFP spatial clustering index. Red crosses and red horizontal lines show mean and SEM. Conditions which do not share a letter are significantly different in Dunn’s multiple comparison test (p< 0.001). Similar results were obtained in three independent experiments. **D**. Proposed graphical summary of our observations for FER-GFP nanoscale dynamics.

**Fig. S7.**
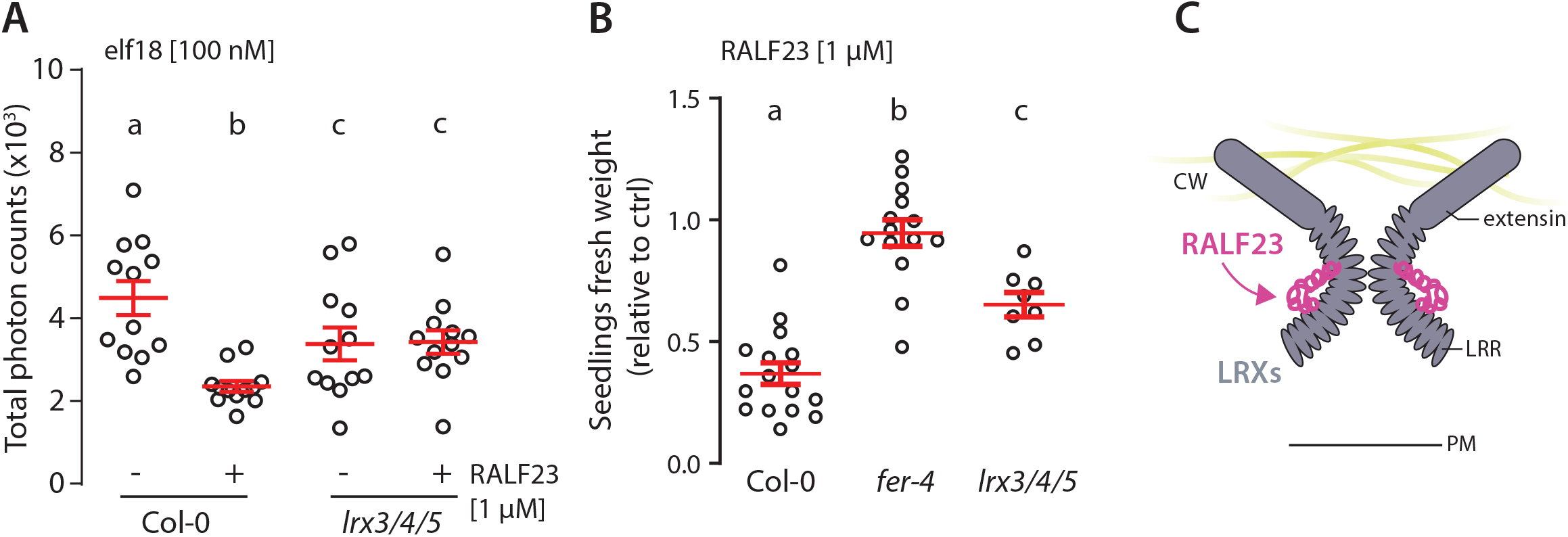
LRX3, LRX4 and LRX5 contribute to RALF23 responsiveness. **A**. ROS production in Col-0 and *lr3/4/5* leaf discs treated with 100 nM elf18 and with or without 1 µM RALF23 co-treatment in 2 mM MES–KOH pH 5.8. Values are means of total photon counts over 40 min. Red crosses and red horizontal lines show mean and SEM, n = 12. Conditions which do not share a letter are significantly different in Dunn’s multiple comparison test (p< 0.0001). **B**. Fresh weight of 12-day-old seedlings grown in the absence (mock) or presence of 1 µM of RALF23 peptide. Fresh weight is expressed as relative to the control mock condition. Similar results were obtained in at least three independent experiments. Conditions which do not share a letter are significantly different in Dunn’s multiple comparison test (p< 0.001). **C**. Proposed graphical representation of LRX3/4/5 as potential receptors for RALF23.

**Fig. S8.**
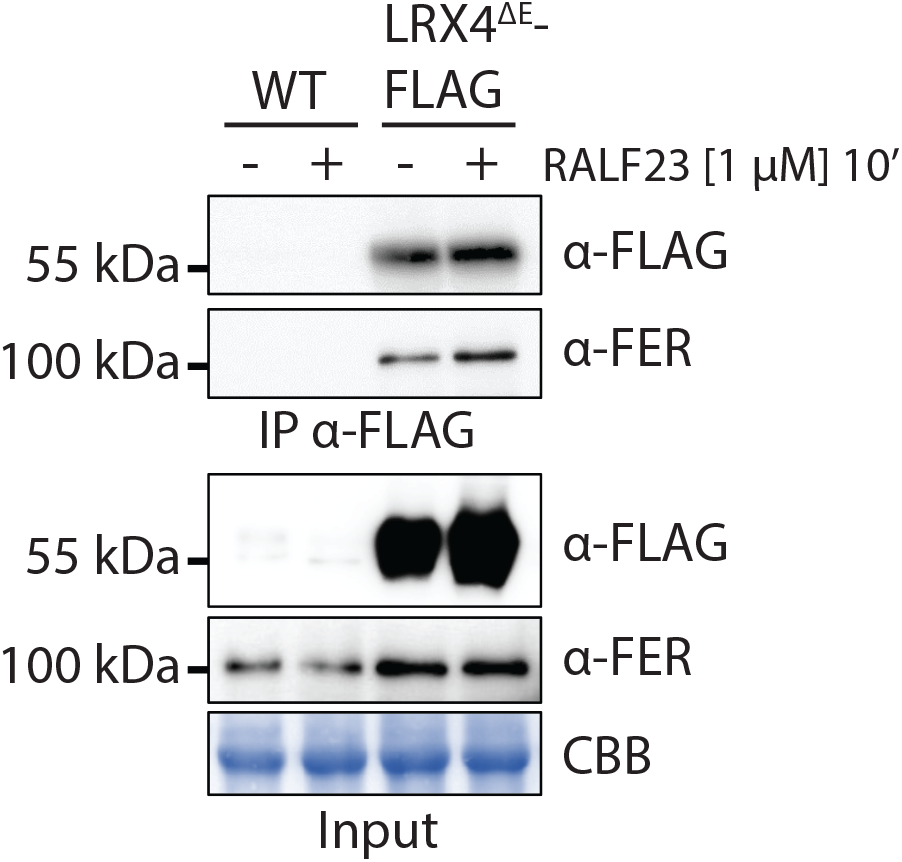
RALF23 does not modulate constitutive association between FER and LRX4. Immunoprecipitation of LRX4^ΔE^-FLAG in Arabidopsis seedlings untreated or treated with 1 µM RALF23 for 10 min. Western blots were probed with α-FLAG or α-FER antibodies. Blot stained with CBB is presented to show equal loading. Similar results were obtained in at least three independent experiments.

**Fig. S9.**
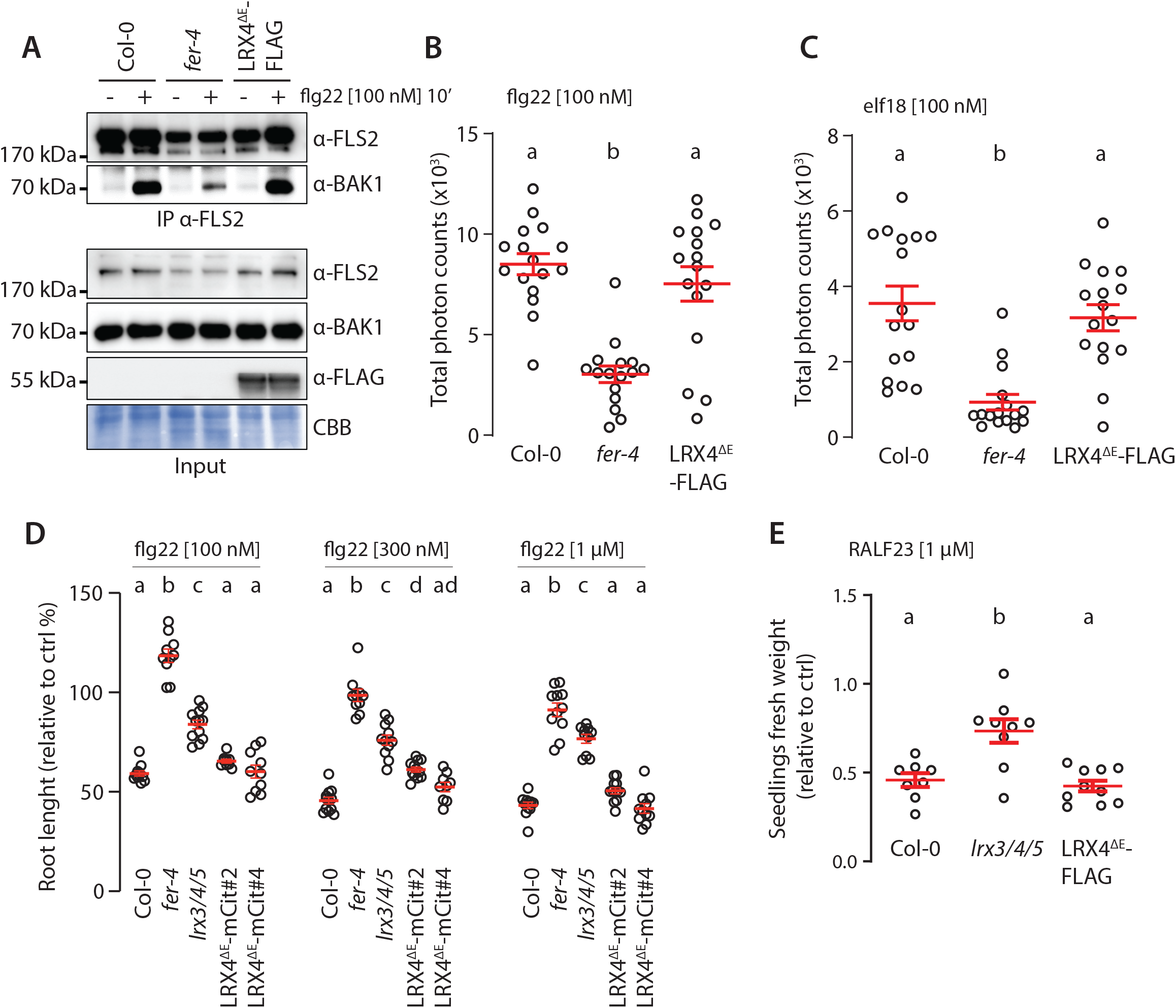
Overexpression of LRX4^ΔE^ does not affect PTI. **A**. flg22-induced FLS2-BAK1 complex formation. Immunoprecipitation of FLS2 in Arabidopsis seedlings either untreated or treated with 100 nM flg22 for 10 min. Blot stained with CBB is presented to show equal loading. Western blots were probed with α-FLS2 and α-BAK1 antibodies. **B-C**. ROS production after elicitation with 100 nM flg22 (B), or 100 nM elf18 (C). Values are means of total photon counts over 40 min. Red crosses and red horizontal lines denote mean and SEM. Conditions which do not share a letter are significantly different in Dunn’s multiple comparison test (p< 0.0001). Similar results were obtained in at least three independent experiments. **D**. Root length of 6-day-old seedlings incubated for 3 days in liquid MS medium with or without indicated concentration of flg22. Red crosses and red horizontal lines denote mean and SEM. Conditions which do not share a letter are significantly different in Brown-Forsythe and Welch ANOVA multiple comparison test (p< 0.001), n= 9-12 seedlings per condition. Similar results were obtained in at least three independent experiments. **E**. Fresh weight of 12-day-old seedlings grown in the absence (mock) or presence of 1 µM of RALF23 peptide. Fresh weight is expressed as relative to the control mock condition. Similar results were obtained in at least three independent experiments. Conditions which do not share a letter are significantly different in Dunn’s multiple comparison test (p< 0.001).

**Fig. S10.**
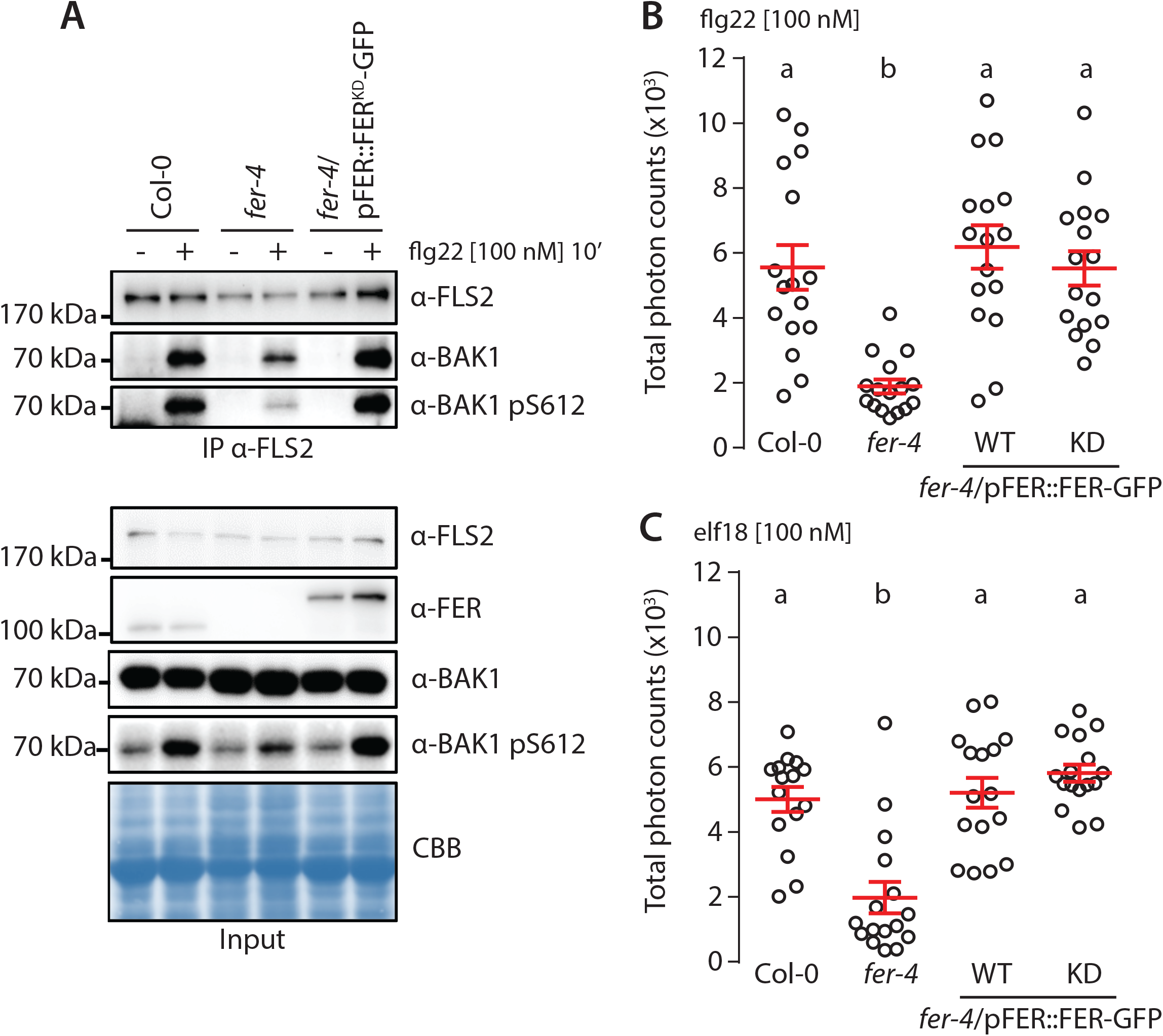
FER kinase activity is dispensable to support PTI signaling. **A**. flg22-induced FLS2-BAK1 complex formation. Immunoprecipitation of FLS2 in Arabidopsis Col-0, *fer-4*, and *fer-4*/pFER::FER^KD^-GFP seedlings that were either untreated or treated with 100 nM flg22 for 10 min. Blot stained with Coomassie brilliant blue (CBB) is presented to show equal loading. Western blots were probed with α-FLS2, α-BAK1, α-BAK1-pS612 or α-FER antibodies. Similar results were obtained in at least three independent experiments. **B-C**. ROS production after elicitation with 100 nM flg22 (**A**), or 100 nM elf18 (**B**). Values are means of total photon counts over 40 min, *n* = 16. Red crosses and red horizontal lines denote mean and SEM, respectively. Conditions which do not share a letter are significantly different in Dunn’s multiple comparison test (p< 0.0001).

**Fig. S11.**
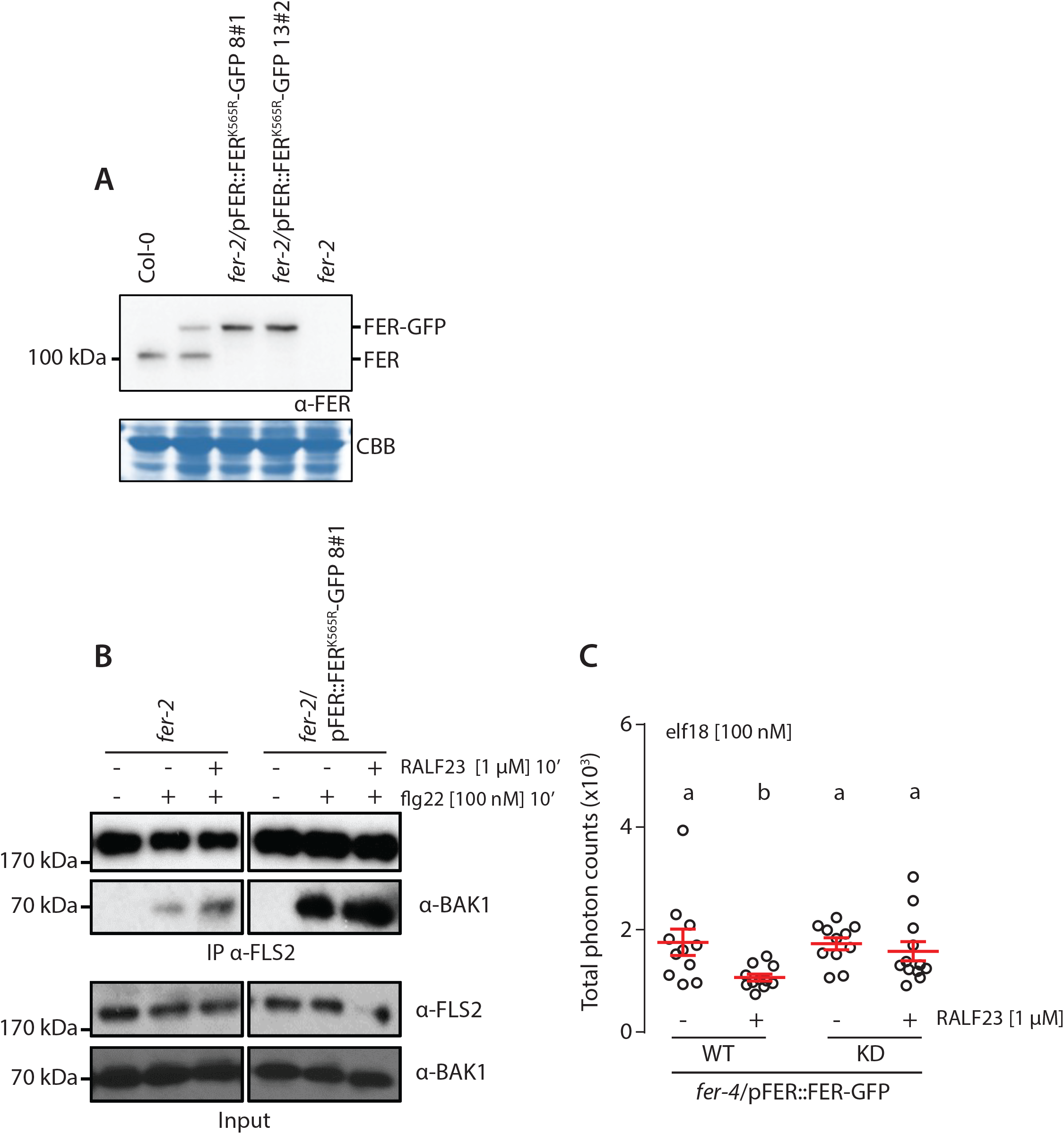
Inhibition of PTI signaling by RALF23 requires FER kinase activity. **A**. Accumulation of endogenous FER and FER^K565R^-GFP detected by western blot. Total proteins were extracted from 2-week-old seedlings. Blot stained with CBB is presented to show loading. **B**. flg22-induced FLS2-BAK1 complex formation. Immunoprecipitation of FLS2 in Arabidopsis *fer-2*, and *fer-2*/pFER::FER^K565R^-GFP seedlings that were either untreated or treated with 100 nM of flg22 and 1 µM RALF23 for 10 min. Western blots were probed with α-FLS2, α-BAK1 antibodies. **C**. ROS production after elicitation with 100 nM elf18, with and without 1 µM RALF23 co-treatment. Values are means of total photon counts over 40 min, n = 10. Red crosses and red horizontal lines denote mean and SEM, respectively. Conditions which do not share a letter are significantly different in Dunn’s multiple comparison test (p< 0.05). Similar results were obtained in at least three independent experiments.

**Fig. S12.**
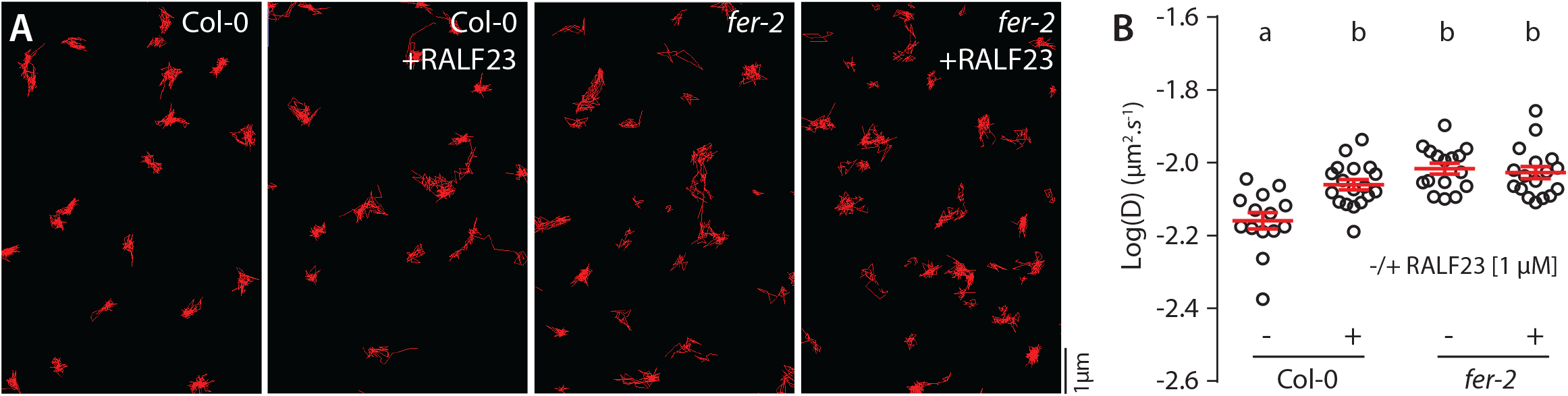
Analysis of FLS2-GFP single particle dynamics upon RALF23 treatment. **A**. Representative super-resolved images of FLS2-GFP single-particles tracked in Col-0 and *fer-2* cotyledon epidermal cells of 5-day-old seedlings with or without 1 µM RALF23 treatment. **B**. and quantification of FLS2-GFP diffusion coefficient (D) in Col-0 and *fer-2* with or without 1 µM RALF23 treatment. Data represent analysis of *ca* 1800 to 2900 single particles observed across 14 to 18 cells. Individual data point represents the mean diffusion coefficient for each cell. Red crosses and red horizontal lines show mean and SEM. Conditions which do not share a letter are significantly different in Dunn’s multiple comparison test (p< 0.0001). Similar results were obtained in at least three independent experiments.

**Fig. S13.**
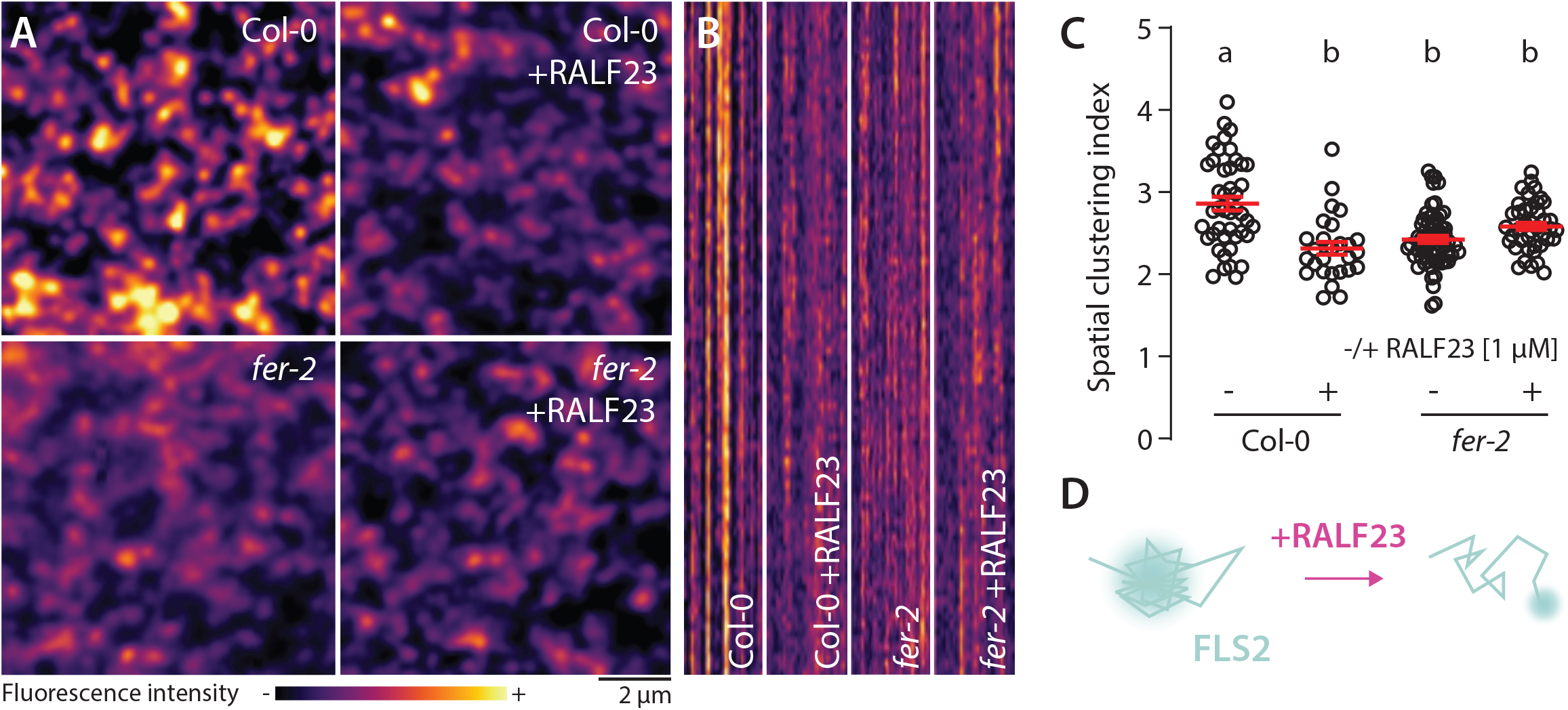
Analysis of FLS2-GFP organization upon RALF23 treatment. **A**. Pictures are maximum projection images (20 TIRFM images obtained at 20 frames per second) of FLS2-GFP in Col-0 and *fer-2* cotyledon epidermal cells with or without 1 µM RALF23 treatment. **B**. Representative kymograph showing lateral organization of FLS2-GFP overtime in Col-0 and *fer-2* cotyledon epidermal cells with or without 1 µM RALF23 treatment. **C**. Quantification of FLS2-GFP spatial clustering index. Red crosses and red horizontal lines show mean and SEM. Conditions which do not share a letter are significantly different in Dunn’s multiple comparison test (p< 0.0001). Similar results were obtained in at least three independent experiments.

**Fig. S14.**
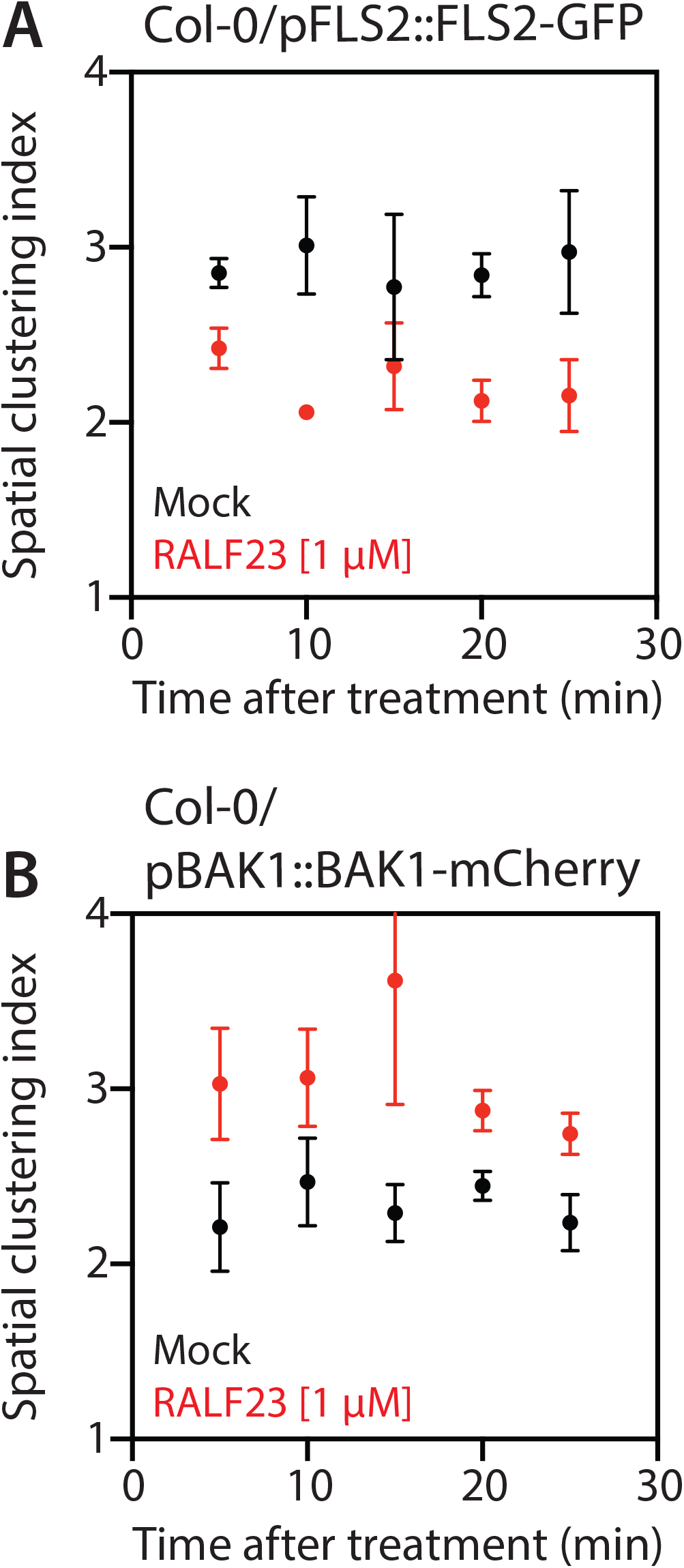
Time-resolved analysis of the spatial clustering index. Data points correspond to the average of the spatial clustering index measured over 5 min windows for FLS2-GFP (**A**) and BAK1-mCherry (**B**) with or without 1 µM RALF23 treatment. The first data point (5 min), corresponds to the value obtained between 2- and 3-min post treatment. Acquisition approximately started 2 min after treatments, the time required for sample mounting.

**Fig. S15.**
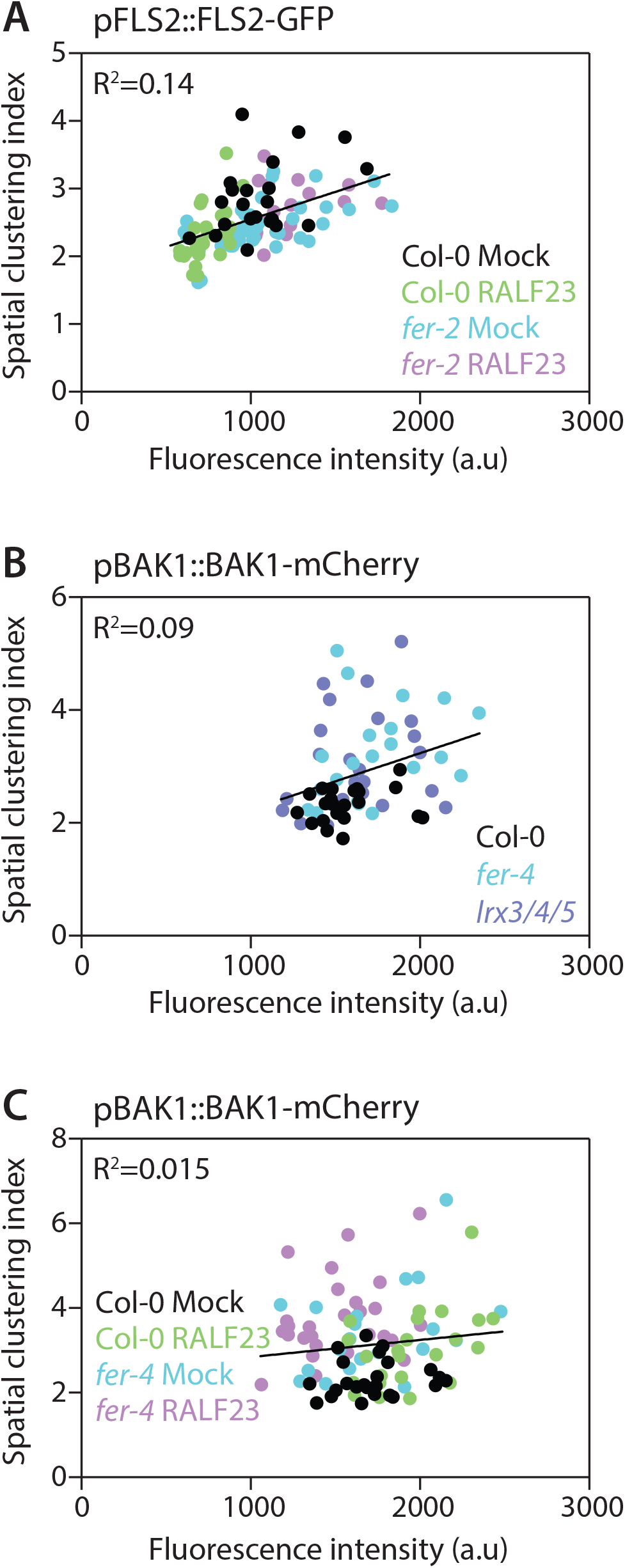
Linear regression analysis of the relationship between the spatial clustering index and fluorescence intensity. Spatial clustering index is calculated as the ratio of the mean of the 5 % highest values to the mean of 5 % lowest values of fluorescence intensity obtain on a plot line. Here, spatial clustering index values were plotted against the mean fluorescence intensity of the corresponding plot line for FLS2-GFP (**A**) and BAK1-mCherry (**B-C**). Linear regression analyses show poor to no correlation between the spatial clustering index and fluorescence intensity (R^2^ ranking from 0.14 and 0.015).

**Movie S1 ¦ TIRFM imaging of FLS2-GFP in Col-0 and *fer-4***.

Representative stream images acquisition of FLS2-GFP particles observed at the surface of 5-day-old cotyledon epidermal cells by TIRF microscopy at 5 frame per second.

**Movie S2 ¦ TIRFM imaging of BAK1-mCherry in Col-0 and *fer-4* with or without RALF23 treatment**.

Representative stream images acquisition of BAK1-mCherry observed at the surface of 5-day-old cotyledon epidermal cells by TIRF microscopy at 2.5 frame per second, in Col-0 (A and B) and *fer-4* (C and D) with (B and D) or without (A and C) 1 μM RALF23 treatment. Scale bar indicates 2 μm.

**Movie S3 ¦ TIRFM imaging of BAK1-mCherry in Col-0 and *lrx3/4/5***.

Representative stream images acquisition of BAK1-mCherry observed at the surface of 5-day-old cotyledon epidermal cells by TIRF microscopy at 2.5 frame per second, in Col-0 (A) and *lrx3/4/5* (B). Scale bar indicates 2 μm.

**Movie S4 ¦ TIRFM imaging of FER-GFP in *fer-4* and *fer-4*;*lrx3/4/5***.

Representative stream images acquisition of FER-GFP observed at the surface of 5-day-old cotyledon epidermal cells by TIRF microscopy at 5 frame per second, in *fer-4* (A) and *fer-4;lrx3/4/5* (B). Scale bar indicates 2 μm.

**Movie S5 ¦ TIRFM imaging of FLS2-GFP in Col-0 with or without RALF23 treatment**.

Representative stream images acquisition of FLS2-GFP particles observed at the surface of 5-day-old cotyledon epidermal cells by TIRF microscopy at 5 frame per second. Scale bar indicates 2 μm.

**Movie S6 ¦ TIRFM imaging of FLS2-GFP in *fer-4* with or without RALF23 treatment**.

Representative stream images acquisition of FLS2-GFP particles observed at the surface of 5-day-old cotyledon epidermal cells by TIRF microscopy at 5 frame per second. Scale bar indicate 2 μm.

**Movie S7 ¦ TIRFM imaging of FLS2-GFP in Col-0 and *fer-2* with or without RALF23 treatment**.

Representative stream images acquisition of FLS2-GFP particles observed at the surface of 5-day-old cotyledon epidermal cells by TIRF microscopy at 20 frame per second. Scale bar indicates 2 μm

## MATERIALS AND METHODS

### Plant materials and growth

*Arabidopsis thaliana* ecotype Columbia (Col-0) was used as WT control. The *fer-4, fer-4/*pFER::FER-GFP (Duan *et al*., 2010), *fer-4/*pFER::FER^KD^-GFP (Chakravorty, Yu and Assmann, 2018), *fer-4/*p35S::FER^ΔMalA^-GFP (Lin *et al*., 2018), Col-0/pFLS2::FLS2-GFP (Göhre *et al*., 2008), *fer-2*/pFLS2::FLS2-GFP (Stegmann *et al*., 2017), *lrx3/4/5*, p35S::LRX4^ΔE^-Citrine and p35S::LRX4^ΔE^-FLAG (Dünser *et al*., 2019) lines were previously published. Col-0/pFLS2::FLS2-GFP (Göhre *et al*., 2008) was crossed with *fer-4* to obtain *fer-4/*pFLS2::FLS2-GFP. Col-0/pBAK1::BAK1-mCherry (Bücherl *et al*., 2013) was crossed with *fer-4* and *lrx3/4/*5 to obtain *fer-*4/pBAK1::BAK1-mCherry and *lrx3/4/5/*pBAK1::BAK1-mCherry. *fer-4/*pFER::FER-GFP was crossed with *lrx3/4/5* to obtain *fer-4/lrx3/4/5;*pFER::FER-GFP. For ROS burst assays, plants were grown in individual pots at 20-21 °C with a 10-h photoperiod in environmentally controlled growth rooms. For seedling-based assays, seeds were surface-sterilized using chlorine gas for 5 h and grown at 22 °C and a 16-h photoperiod on Murashige and Skoog (MS) medium supplemented with vitamins, 1 % sucrose and 0.8 % agar.

### Synthetic peptides and chemicals

The flg22, elf18, and RALF23 peptides were synthesized by EZBiolab (United States) with a purity of >95 %. All peptides were dissolved in sterile purified water.

### ROS burst measurement

ROS burst measurements were performed as previously documented (Kadota *et al*., 2014). At least eight leaf discs (4 mm in diameter) per individual genotype were collected in 96-well plates containing sterile water and incubated overnight. The next day, the water was replaced by a solution containing 17 μg/mL luminol (Sigma Aldrich), 20 μg/mL horseradish peroxidase (HRP, Sigma Aldrich) and the peptides in the appropriate concentration. Luminescence was measured for the indicated time period using a charge-coupled device camera (Photek Ltd., East Sussex UK). The effect of RALF23 on elf18-triggered ROS production was performed as previously described (Stegmann *et al*., 2017). Eight to ten leaf discs per treatment and/or genotype were collected in 96-well plates containing water and incubated overnight. The following day, the water was replaced by 75 µL of 2 mM MES-KOH pH 5.8 to mimic the apoplastic pH. Leaf discs were incubated further for 4-5 h before adding 75 μL of a solution containing 40 μg/mL HRP, 1 μM L-O12 (Wako Chemicals, Germany) and 2X elicitor RALF peptide solution (final concentration 20 μg/mL HRP, 0.5 µM L-O12, 1x elicitors). ROS production is displayed as the integration of total photon counts.

### Root growth inhibition assay

Three-day-old Col-0, *fer-4, lrx3/4/5* and 35S::LRR4-Cit seedlings (n = 9-12) were transferred for additional 3 days to 3 mL liquid ½ MS medium containing different concentrations (100 nM, 300 nM or 1 µM) of flg22 or the appropriate amount of solvent. The seedlings were then placed on solid MS plates before scanning. Root length was measured using ImageJ.

### Live cell imaging

For confocal microscopy and TIRF microscopy experiments, surface-sterilized seeds were individually placed in line on square Petri dishes containing 1/2 MS 1 % sucrose, 0.8 % phytoagar, stratified 2 d in the dark at 4 °C, then placed in a growth chamber at 22 °C and a 16-h photoperiod for 5 d. Seedlings were mounted between a glass slide and a coverslip in liquid 1/2 MS, 1 % sucrose medium. To test the effect of RALF23 on FLS2-GFP dynamics and nanodomain organization, seedlings were pre-incubated in 2 mM MES-KOH pH 5.8 for 3 to 4h prior treatment. Seedlings were image 2-30 min after treatment.

### Confocal laser scanning microscopy (CLSM)

Confocal microscopy was performed using a Leica SP5 CLSM system (Leica, Wetzlar, Germany) equipped with Argon, DPSS, He-Ne lasers, hybrid detectors and using a 63X 1.2 NA oil immersion objective. GFP was excited using 488 nm argon laser and emission wavelengths were collected between 495 and 550 nm. mCherry was excited using 561 nm He/Ne laser and emission wavelengths were collected between 570 and 640 nm. Propidium iodide was imaged using 488 nm and 500-550 nm excitation and emission wavelengths, respectively. In order to obtain quantitative data, experiments were performed using strictly identical confocal acquisition parameters (*e*.*g*. laser power, gain, zoom factor, resolution, and emission wavelengths reception), with detector settings optimized for low background and no pixel saturation. Pseudo-color images were obtained using look-up-table (LUT) of Fiji software (Schindelin *et al*., 2012).

### Total Internal Reflection Fluorescence (TIRF) microscopy

TIRF microscopy was performed using an inverted Leica GSD equipped with a 160x objective (NA = 1.43, oil immersion), and an Andor iXon Ultra 897 EMCCD camera. Images were acquired by illuminating samples with a 488 nm solid state diode laser set at 15 mW, using a cube filter with an excitation filter 488/10 and an emission filter 535/50 for FLS2-GFP and FER-GFP. optimum critical angle was determined as giving the best signal-to-noise. Images time series were recorded at 20 frames per second (50 ms exposure time) for Figure S11 and Figure S3; 5 frames per second for Figure 1 and Figure S6. To observe BAK1-mCherry we could only use a 532 nm solid state diode laser (*c*.*a* 40% of maximum excitation for mCherry), using a cube filter with an excitation filter 532/10 and an emission filter 600/100. To obtain a sufficient signal to noise ratio images time series were recorded at 2.5 frames per second (Figure 1, 2 and 5). Due to apparent high mobility of BAK1 and relatively slow acquisition rate we couldn’t asses with confidence the identity of fluorescent particles from one time frame to another and therefore did not perform particle tracking analysis of BAK1-mCherry.

### Single particle tracking analysis

To analyse single particle tracking experiments, we used the plugin TrackMate 2.7.4 (Tinevez *et al*., 2017) in Fiji (Schindelin *et al*., 2012). Single particles were segmented frame-by-frame by applying a LoG (Laplacian of Gaussian) filter and estimated particle size of 0.4 μm. Individual single particle were localized with sub-pixel resolution using a built-in quadratic fitting scheme. Then, single particle trajectories were reconstructed using a simple linear assignment problem (Jaqaman *et al*., 2008) with a maximal linking distance of 0.4 μm and without gap-closing. Only tracks with at least ten successive points (tracked for 500 ms) were selected for further analysis. Diffusion coefficients of individual particles were determined using TraJClassifier (Wagner *et al*., 2017). For each particle, the slope of the first four time points of their mean square displacement (MSD) plot was used to calculate their diffusion coefficient according to the following equation: MSD= (x-x_0_)^2^+(y-y_0_)^2^ and D=MSD/4*t*, where x0 and y0 are the initial coordinates, and x and y are the coordinates at any given time, and *t* is the time frame.

### Quantification of spatial clustering index

Genotype and/or treatment dependent variation in fluorescence intensity of FLS2-GFP and fluorescence pattern of FLS2-GFP and BAK1-mCherry compromised the use of a unique set of parameters to compute nanodomain size and density across the different experiments. To uniformly quantify differences in membrane organization of both FLS2 and BAK1 across all experiments we used the spatial clustering index which was shown to be largely insensitive to variation in fluorescence intensity (Gronnier *et al*., 2017). Quantifications were performed as previously described (Gronnier *et al*., 2017). Briefly, fluorescence intensity was plotted along an 8 µm long line on maximum projection TIRFM images, three plots were randomly recorded per cell and, at least 8 cells per condition per experiment were analysed. For each line plot, the spatial clustering index was calculated by dividing the mean of the 5 % highest values by the mean of 5 % lowest values. Because the absence of correlation between fluorescence intensity and spatial clustering index was assessed on confocal microscopy images and for a single protein (Gronnier *et al*., 2017), we decided to test if this was also the case in our experimental conditions. Indeed, we consistently observed poor to no correlation between variation in fluorescence intensity and values of spatial clustering index (Sup Fig. 15).

### Co-immunoprecipitation experiments

Twenty to thirty seedlings per plate were grown in wells of a 6-well plate for 2 weeks, transferred to 2 mM MES-KOH, pH 5.8 and incubated overnight. The next day, flg22, (final concentration 100 nM) and/or RALF23 (final concentration 1 µM) were added and incubated for 10 min. Seedlings were then frozen in liquid N2 and subjected to protein extraction. To analyse FLS2-BAK1 receptor complex formation, proteins were isolated in 50 mM Tris-HCl pH 7.5, 150 mM NaCl, 10 % glycerol, 5 mM dithiothreitol, 1 % protease inhibitor cocktail (Sigma Aldrich), 2 mM Na2MoO4, 2.5 mM NaF, 1.5 mM activated Na3VO4, 1 mM phenylmethanesulfonyl fluoride and 0.5 % IGEPAL. For immunoprecipitations, α-rabbit Trueblot agarose beads (eBioscience) coupled with α-FLS2 antibodies (Chinchilla *et al*., 2007) or GFP-Trap agarose beads (ChromoTek) were used and incubated with the crude extract for 3-4 h at 4 °C. Subsequently, beads were washed 3 times with wash buffer (50 mM Tris-HCl pH 7.5, 150 mM NaCl, 1 mM phenylmethanesulfonyl fluoride, 0,1 % IGEPAL) before adding Laemmli sample buffer and incubating for 10 min at 95 °C. Analysis was carried out by SDS-PAGE and immunoblotting. To test the association between Flag-LRX4 and FER, total protein from 60-90 seedlings per treatment per genotype was extracted as previously described. For immunoprecipitations, M2 anti-Flag affinity gel (Sigma A2220-5ML) was used and incubated with the crude extract for 2-3 h at 4 °C. Subsequently, beads were washed 3 times with wash buffer (50 mM Tris-HCl pH 7.5, 150 mM NaCl, 1 mM phenylmethanesulfonyl fluoride, 0,1 % IGEPAL) before adding Laemmli sample buffer and incubating for 10 min at 95 °C. Analysis was carried out by SDS-PAGE and immunoblotting.

### Immunoblotting

Protein samples were separated in 10 % bisacrylamide gels at 150 V for approximately 2 h and transferred into activated PVDF membranes at 100 V for 90 min. Immunoblotting was performed with antibodies diluted in blocking solution (5 % fat-free milk in TBS with 0.1 % (v/v) Tween-20). Antibodies used in this study: α-BAK1 (1:5 000; (Roux *et al*., 2011); α-FLS2 (1:1000; (Chinchilla et al., 2007); α-FER (1:2000; (Xiao *et al*., 2019), α-BAK1 pS612 (1:3000; (Perraki *et al*., 2018)), α-FLAG-HRP (Sigma Aldrich, A8592, dilution 1:4000); α -GFP (sc-9996, Santa Cruz, used at 1:5000). Blots were developed with Pierce ECL/ ECL Femto Western Blotting Substrate (Thermo Scientific). The following secondary antibodies were used: anti-rabbit IgG-HRP Trueblot (Rockland, 18-8816-31, dilution 1:10000) for detection of FLS2-BAK1 co-immunoprecipitation or anti-rabbit IgG (whole molecule)–HRP (A0545, Sigma, dilution 1:10000) for all other western blots.

### Statistical analysis

Statistical analyses were carried out using Prism 6.0 software (GraphPad). As mentioned in the figure legend, statistical significances were assessed using non-parametric Kruskal-Wallis bilateral tests combined with post-hoc Dunn’s multiple pairwise comparisons, or using a two-way non-parametric student’s t test Mann-Whitney test.

### Accession numbers

FER (AT3G51550), LRX3 (AT4G13340), LRX4 (AT3G24480), LRX5 (AT4G18670), RALF23 (AT3G16570), FLS2 (AT5G46330), BAK1 (AT4G33430).

